# CRISPR/Cas9-based depletion of 16S ribosomal RNA improves library complexity of single-cell RNA-sequencing

**DOI:** 10.1101/2023.05.25.542286

**Authors:** Kuang-Tse Wang, Carolyn E. Adler

## Abstract

**Background:** Single-cell RNA-sequencing (scRNA-seq) relies on PCR amplification to retrieve information from vanishingly small amounts of starting material. To selectively enrich mRNA from abundant non-polyadenylated transcripts, poly(A) selection is a key step during library preparation. However, some transcripts, such as mitochondrial genes, can escape this elimination and overwhelm libraries. Often, these transcripts are removed *in silico*, but whether physical depletion improves detection of rare transcripts in single cells is unclear.

**Results:** We find that a single 16S ribosomal RNA is widely enriched in planarian scRNA-seq datasets, independent of the library preparation method. To deplete this transcript from scRNA-seq libraries, we design 30 single-guide RNAs spanning its length. To evaluate the effects of depletion, we perform a side-by-side comparison of the effects of eliminating the 16S transcript and find a substantial increase in the number of genes detected per cell, coupled with virtually complete loss of the 16S RNA. Moreover, we systematically determine that library complexity increases with a limited number of PCR cycles following CRISPR treatment. When compared to *in silico* depletion of 16S, physically removing it reduces dropout rates, retrieves more clusters, and reveals more differentially-expressed genes.

**Conclusions:** Our results show that abundant transcripts reduce the retrieval of informative transcripts in scRNA-seq and distort the analysis. Physical removal of these contaminants enables the detection of rare transcripts at lower sequencing depth, and also outperforms *in silico* depletion. Importantly, this method can be easily customized to deplete any abundant transcript from scRNA-seq libraries.

## Background

Recent advances in single-cell transcriptomics (scRNA-seq) have greatly facilitated the exploration of cell type complexity. By capturing and barcoding mRNA on a cell-by-cell basis, scRNA-seq enables researchers to identify different cell types in a wide range of species. However, an important aspect of scRNA-seq is quality control in the analysis, which involves removing potential confounding factors such as sequencing depth and the percentage of mitochondrial reads (1,2). While these factors can be normalized *in silico*, it is optimal to minimize unwanted transcripts during library preparation to improve efficiency and more accurately probe cell complexity.

To maximize retrieval of protein-coding transcripts, most scRNA-seq methods capture mRNA by poly(A) enrichment (3,4). Ideally, poly(T) primers selectively bind to polyadenylated protein-coding transcripts to enable reverse transcription. This approach effectively eliminates ribosomal RNA, which comprises >80% of total RNA. Despite its utility, poly(A) enrichment also prevents the capture of some informative RNA species, including histone mRNA, miRNA, and enhancer RNAs (5). In addition, unwanted transcripts such as mitochondrial RNA leak out of damaged cells and are carried through into cDNA libraries (2). These contaminants often appear as ambient RNA, a computational challenge for cell calling algorithms that can introduce bias into normalization (6,7). Due to these issues, poly(A) enrichment can distort the global transcriptional landscape.

Alternative ribodepletion methods have recently been developed for single-cell transcriptomics. For example, VASA-seq generates cDNA by hexamer primers with a T7 handle to enable subsequent RNA amplification by in vitro transcription. Ribosomal RNAs will then be removed from amplified RNA by rDNA probes and RNAse-H-mediated digestion (8). RamDA-seq uses specific hexamers that are not present in rRNA to exclude it from downstream steps (9,10). However, these methods require new chemistry and reagents, raising the entry barrier for in-house setup.

Depletion of Abundant Sequences by Hybridization (DASH) is a CRISPR/Cas9-based method to remove unwanted DNA sequences from any DNA library (11). With the customized design of single guide RNA (sgRNA), Cas9 can precisely remove unwanted cDNA sequences, and further enhance the detection of rare transcripts. DASH has not only been shown to efficiently deplete unwanted sequences in 16S sequencing and bulk RNA-seq (12,13) but has been adapted to single-cell transcriptome methods, including scCLEAN, Smart-seq-total and MATQ-seq (14–17). However, whether this depletion impacts library complexity or other metrics of sequencing quality has yet to be tested systematically.

Planarian mitochondrial 16S rRNA sequences are known to make up ∼30% of bulk RNA-seq libraries generated by poly(A) enrichment (18). In scRNA-seq experiments, most studies remove 16S from the analysis, but to what extent 16S rRNA contaminates scRNA-seq remains elusive (19,20). Here, we reanalyzed published datasets and found that unique molecular identifiers (UMIs) mapping to 16S constitute approximately 5-74% of sequencing reads regardless of the single-cell library generation strategy. We adapted DASH, which leverages CRISPR to remove unwanted sequences, to the 10X Chromium protocol to deplete mitochondrial 16S cDNA from planarian sequencing libraries (18). We sequenced the same library with and without DASH treatment and carried out a side-by-side analysis to determine the impact of DASH treatment on overall scRNA-seq performance. We show that our protocol specifically depletes more than 90% of 16S UMIs from both cells and ambient RNA. This depletion increases the number of genes and non-16S UMIs detected per cell which improves the downstream analysis. We conclude that DASH can enhance library complexity, boosting the information retrieved from scRNA-seq experiments, with significant economic benefits. Importantly, this technique can be easily customized to deplete any abundant transcript after single-cell libraries are generated.

## Results

### Mitochondrial 16S transcript dominates reads in planarian scRNA-seq datasets

Planarians were one of the first animal models to be profiled at the whole organismal level using scRNA-seq. Because several different sequencing strategies have been applied to planarians, we first asked which method has the highest library complexity and the least 16S rRNA contamination. Thus, we collected 13 datasets from 7 studies using Drop-seq (19,21), Smart-seq2 (22), Split-Seq (23), Split-Seq with ACME fixation (24), and 10X Chromium (20,25) (Figure 1). All methods use poly(T) primers to capture transcripts except for Split-Seq, which uses a mixture of random hexamers and poly(T) primers. To standardize these comparisons across libraries, we used the same pipeline to pre-process the barcodes and UMI tagging, align the reads to the genome (including the mitogenome), and generate quality metrics (26,27).

**Figure 1.**
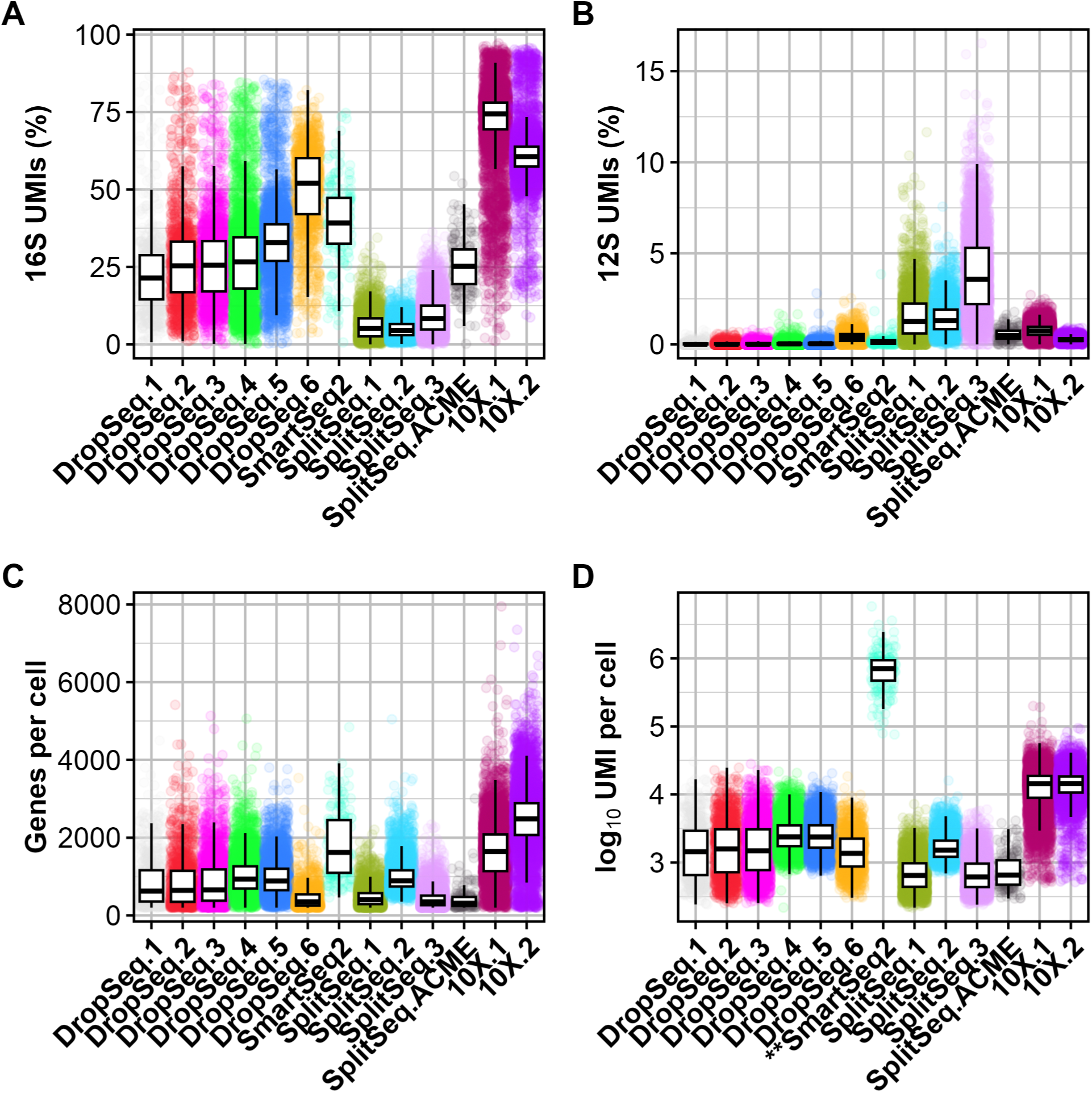
Planarian 16S rRNA is highly enriched in various scRNA-seq platforms. **A-B,** Boxplots show the percentage of 16S UMIs (**A**) and 12S UMIs (**B**) across different published scRNA-seq datasets. **C-D,** Boxplots show the number of genes (**C**) and UMI (**D**) per cell. Note, SmartSeq2 does not implement UMI. The values graphed indicate total read counts for SmartSeq2. The boxes show the 25th, 50th, and 75 quantiles and the whiskers show 1.5 times interquartile range. DropSeq.1-5: “head” datasets (19); DropSeq.6:gfp(RNAi) (21); SmartSeq2 (22); SplitSeq.1-3: 0 days post-amputation (dpa), 1dpa and 2dpa (23); SplitSeq.ACME (24); 10X.1: post-pharyngeal wound fragments, 0h (25); 10X.2 (20).

To assess the abundance and quality of these different sequencing strategies, we applied several metrics. First, we analyzed the percentage of reads that mapped to the 16S transcript. Libraries made with 10X Chromium had the highest percentage of UMIs mapping to this transcript (61% and 74% on average) (Figure 1A). By contrast, SplitSeq libraries were the lowest (ranging from 5% to 8%). DropSeq and ACME-based strategies ranged from 21 to 52%. The prevalence of 16S was unique because another mitochondrial rRNA, 12S, was consistently low across all datasets (Figure 1B). These findings showed that in various scRNA-seq methods, the single 16S transcript accounted for a significant fraction of UMIs. Although Split-Seq appears to have the lowest percentage of UMIs mapping to the 16S, these libraries also had the highest number of unmappable reads (Supplemental Table 1). Next, we examined library complexity by assessing the number of genes and UMIs per cell. While 10X Chromium retained the highest levels of 16S UMIs as compared to other tested library methodologies, it also yielded the greatest library complexity, as assessed by the genes and UMIs detected per cell (Figure 1C-D). In conclusion, these results show that 16S contamination is a severe problem in scRNA-seq preparation for planarians.

Because all of the scRNAseq library strategies rely on poly(A) enrichment, we hypothesized that 16S may be retained because of either polyadenylation or an internal polyA stretch (28,29). To distinguish between these two possibilities, we analyzed the coverage of reads across the 16S locus in one of our datasets using 10X Chromium. We found that the reads from a 10X dataset were strongly skewed to the 3’ end (Supplemental Figure 1A), similar to other protein-coding genes, suggesting polyadenylation of 16S (Supplemental Figure 1B). We also identified poly(A) stretches close to the middle of the 16S sequence (Supplemental Figure 1B). Although these data did not provide a clear reason for the high abundance of 16S rRNA, we sought an alternative method to exclude 16S rRNA for scRNA-seq preparation.

### DASH effectively depletes 16S cDNA in scRNA-seq libraries

Due to the widespread use of the 10X Chromium scRNA-seq platform, and because it yields the highest library complexity in planarians, we sought to optimize the 10X protocol by selectively removing the 16S sequence. During single-cell RNAseq library preparation, reverse transcription occurs immediately after cell lysis, concomitant with barcoding, so any depletion must happen during or after this step (4). Ribodepletion methods that remove unwanted RNA, such as RNAse-H mediated digestion, are difficult to integrate with 10X scRNA-seq because of the barcoding that occurs concomitant with cDNA generation. Depletion of Abundant Sequences by Hybridization (DASH) is a method that effectively depletes cDNA in a sequence-specific manner, making it a promising method to eliminate the 16S transcript after barcoding (11). Therefore, we designed 30 non-overlapping single guide RNAs (sgRNAs) tiling the entire 16S transcript. To eliminate this transcript, we reasoned that degrading 16S as early as possible after cDNA conversion would be optimal to preserve the transcriptional repertoire. We integrated DASH into the 10X Chromium workflow by only performing 10 PCR cycles after cDNA conversion, then incubating the cDNA with pooled sgRNAs complexed with Cas9. After CRISPR/Cas9 degradation, we further amplified cDNA with 10 additional PCR cycles, followed by the standard end repairing and indexing steps (Figure 2A).

**Figure 2.**
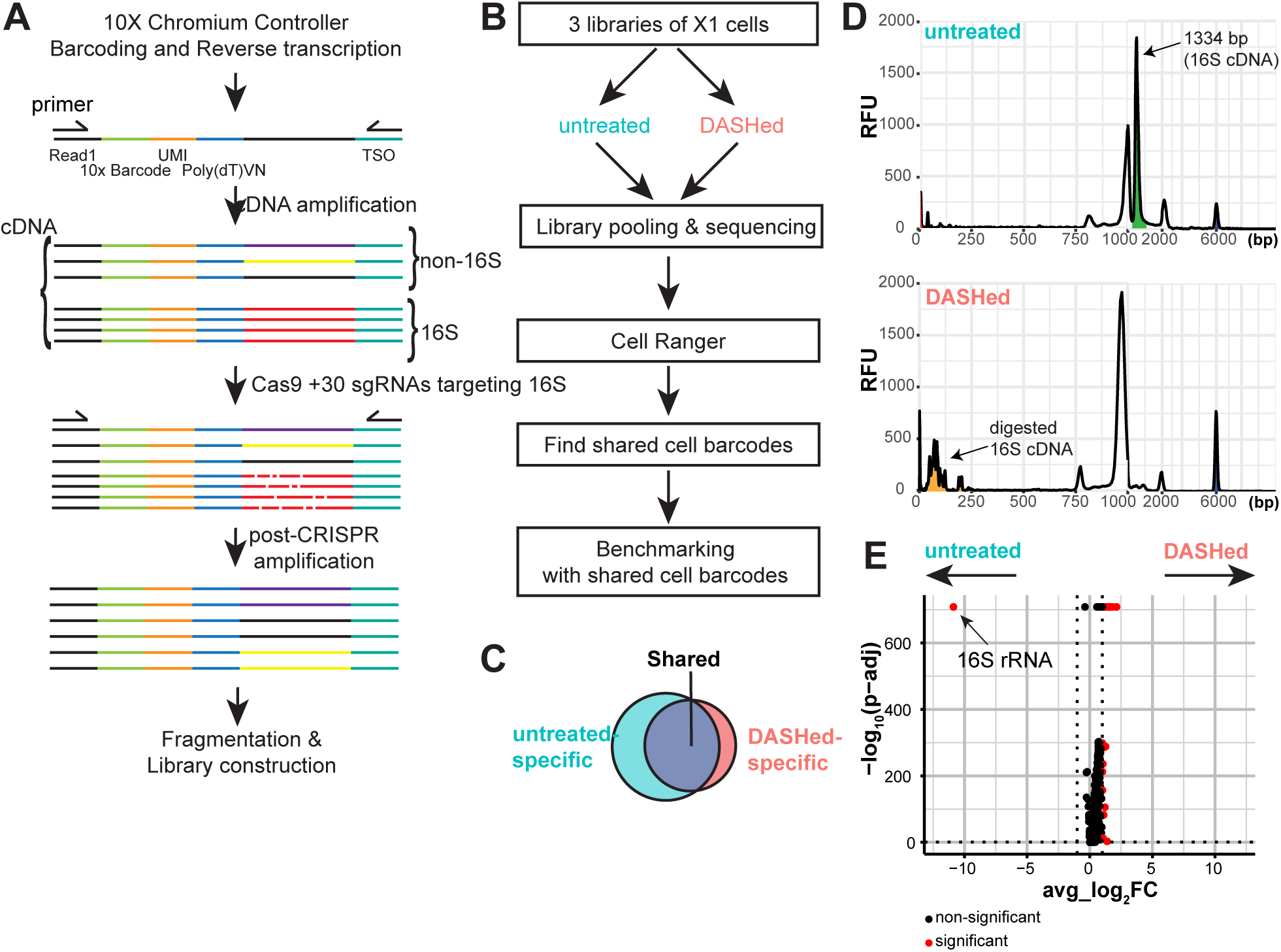
Workflow and depletion of 16S sequence with DASH. **A,** Schematic of DASH protocol. Probes containing poly(T)VN, where V is any base except T, and N is any base, are used for poly(A) enrichment. cDNA is reverse-transcribed in the 10X Chromium Controller and amplified by cDNA primers, Read1 and template switching oligo (TSO). After generation of cDNA, 16S cDNA is depleted by incubation with Cas9 and sgRNAs targeting 16S sequence, followed by post-CRISPR amplification with the same cDNA primers. The re-amplified cDNA is then fragmented and indexed for subsequent sequencing. **B,** Schematic of experimental design to benchmark the performance of DASH. Three biological replicates of stem cells (X1 cells) are sorted and processed for cDNA preparation. cDNA is then split into “untreated” and “DASHed” libraries. Sequencing reads are processed by CellRanger and then the cell barcodes recovered from both groups are used for downstream analysis. **C,** Venn diagram of cell barcodes from untreated and DASHed libraries. The shared cell barcodes are used to assess library quality. **D,** Fragment analysis of untreated cDNA (**A**) and DASHed cDNA (**B**). x-axis shows fragment size in base pairs (bp), and y-axis shows relative fluorescence units (RFU). Red peak marks the lower marker (1bp) and blue peak marks the upper marker (6000bp). **E,** Volcano plot of differential expression analysis of DASHed versus untreated datasets. Data includes 3 biological replicates of scRNA-seq libraries combined and analyzed as pseudobulk samples. x-axis is the average log2 fold change across all cells. y-axis is -log10 of adjusted p-value of Wilcoxon Rank Sum test based on Bonferroni correction. The cutoff for significant genes is adjusted p-value < 0.05 and absolute average log2 fold change > 1.

To test whether DASH could deplete 16S sequences from cDNA libraries, we first generated three 10X 3’ scRNA-seq cDNA libraries from planarian stem cells. Next, we split the cDNA libraries and generated DASH-treated (‘DASHed’) libraries for each biological replicate (Figure 2B). By sequencing the same biological libraries before and after DASH treatment, we could benchmark the impact of DASH treatment at the single-cell level by analyzing shared cell barcodes (Figure 2C).

To assess the efficiency of 16S depletion, we compared untreated and DASHed libraries on the Bioanalyzer. Fragment analysis showed a strong peak around 1334bp in the untreated cDNA that was absent in the DASHed cDNA, while the rest of the profile remained comparable (Figure 2D). Planarian 16S rRNA is predicted to be ∼900bp long (18,30). Even with the addition of 112bp of primers from the 10X library (82bp, poly(dT) primer + 30bp, TSO primer), this does not account for the discrepancy in length, raising the possibility that this peak may not represent the true 16S transcript. To verify that we depleted the 16S transcript, we compiled 3 scRNA-seq replicate libraries before and after DASH treatment into a pseudobulk plot. Indeed, differential expression analysis showed that the 16S rRNA was the only downregulated gene in DASHed datasets (Figure 2E). This specific downregulation suggests that the highest peak in the untreated library may represent 16S cDNA and that it can be efficiently depleted by DASH.

### Overrepresentation of 16S UMIs causes aberrant cell calling

Cell calling algorithms rely on the assumption that cells have significantly more UMIs than empty droplets and ambient RNA, so they can distinguish cells from non-cells by total UMIs associated with each cell barcode (4). We hypothesized that the high prevalence of 16S rRNA could potentially interfere with cell calling by inappropriately ‘calling’ cells that in reality are ambient RNA. To evaluate the effect of DASH treatment on cell calling, we obtained lists of cell barcodes from untreated and DASHed libraries generated by Cell Ranger for each of the 3 biological replicates (Figure 2C). The majority of cell barcodes were shared between untreated and DASHed datasets for each replicate (Supplemental Figure 2A). Across all 3 replicates, the untreated datasets consistently had more cell barcodes called by Cell Ranger as cells (ranging from 9% to 20% of the total) (Supplemental Figure 2A). Of the barcodes that were unique to the untreated libraries, the vast majority (>90%) mapped to 16S UMIs (Supplemental Figure 2B). Barcodes shared by both untreated and DASHed libraries had 58-60% of 16S UMIs in untreated datasets, which is comparable to previously published datasets (Supplemental Figure 2B; Figure 1A). By contrast, the barcodes that were specific to the DASHed libraries had very low levels of 16S UMIs (<0.2%), indicating that the depletion was thorough (Figure 2E). Together, these results suggest that 16S comprises a significant proportion of total UMIs in both cell and ambient RNA and misleads the cell calling process.

### DASH depletes 16S rRNA and increases mRNA and gene recovery

In general, mitochondrial and ribosomal sequences are computationally removed from downstream analysis (19,20). Because depletion with DASH is so specific, it may achieve the same effect as this computational processing step. Alternatively, the physical depletion of 16S at an early step in library generation could recover more sequencing information than computational removal. To assess the efficacy of 16S depletion on library complexity, we normalized untreated and DASHed scRNA-seq datasets to 100 million reads by downsampling and examined metrics of library quality. The 16S UMIs dropped from 60% to less than 0.2% in DASHed datasets, suggesting that the 16S cDNA was efficiently depleted (Figure 3A). Moreover, the total number of genes detected per cell showed an increase ranging from 27% to 40% (Figure 3B), and the non-16S UMIs (mostly mapped to protein-coding transcripts) increased from 46% to 70% in DASHed datasets as compared to untreated libraries (Figure 3C). These findings suggest that DASH treatment can enrich library complexity further than computational depletion alone (Figure 3C-D).

**Figure 3.**
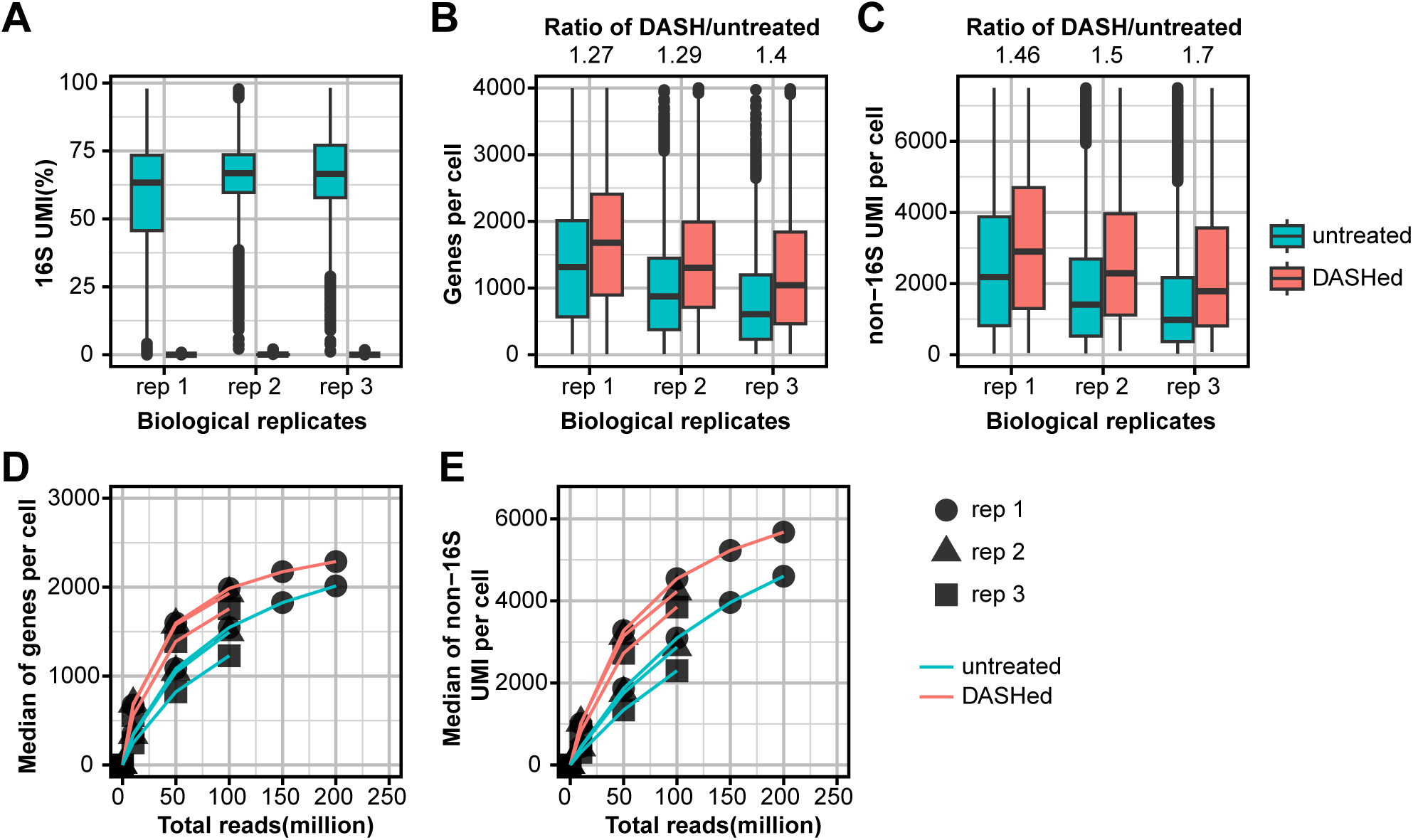
DASH depletes 16S UMIs and enhances library complexity. **A-C,** Boxplots show the percentage of 16S UMI (**A**), numbers of genes (**B**) and non-16S UMIs (**C**) per cell in three replicates before and after DASH treatment. The medians of ratio in DASHed versus untreated are shown on the top. All samples from 3 biological replicates (rep) contain 100 million reads. **D-E,** Rarefaction analysis of library complexity comparing the libraries between untreated and DASHed, shown are the medians of genes (**D**) and non-16S UMI per cell (**E**). Each dot represents a downsampled replicate with indicated total reads.

To investigate whether the loss of complexity in the untreated library could be overcome by simply increasing sequencing reads, we performed a rarefaction analysis. This analysis assesses library complexity by measuring how diversity increases with sequencing reads until it reaches saturation. We downsampled the datasets to 10, 50, and 100 million reads for replicates 2 and 3, and 150 and 200 million reads for replicate 1, and examined the library quality metrics of shared cell barcodes. We found that the number of genes detected per cell and non-16S UMIs were consistently higher in DASHed libraries, even at the lowest read depths (Figure 3D-E). These results indicate that the depletion of 16S rRNA consistently enhances library complexity given the same amount of reads, beyond what can be achieved by computational ribodepletion.

### Post-CRISPR amplification is necessary and improves library complexity

In scRNA-seq methods, the number of PCR cycles to amplify cDNA is adjusted based on the estimated cell numbers, tested empirically by the 10X manufacturer (4). Over-or under-amplification leads to a decrease in library complexity. We asked if the post-CRISPR amplification is necessary and if so, what are the optimal PCR cycles that maximize library diversity without changing the overall fragment distribution. We sequenced replicate 1 without any re-amplification (0 cycles), or with 5, 10, and 15 PCR cycles, and assessed library complexity by downsampling into the same total reads. We excluded the cDNA amplified with 15 cycles from further sequencing because the cDNA trace changed drastically as compared to 5 and 10 cycles (Supplemental Figure 3). After sequencing, the number of UMIs and genes per cell increased with more PCR cycles (Figure 4A-B). Similarly, the rarefaction analysis also showed that library complexity was consistently higher in the 10-cycle condition (Figure 4C-D). This finding suggests that post-CRISPR amplification is necessary for improving library complexity, with an optimal cycle number of 10, at least for these samples.

**Figure 4.**
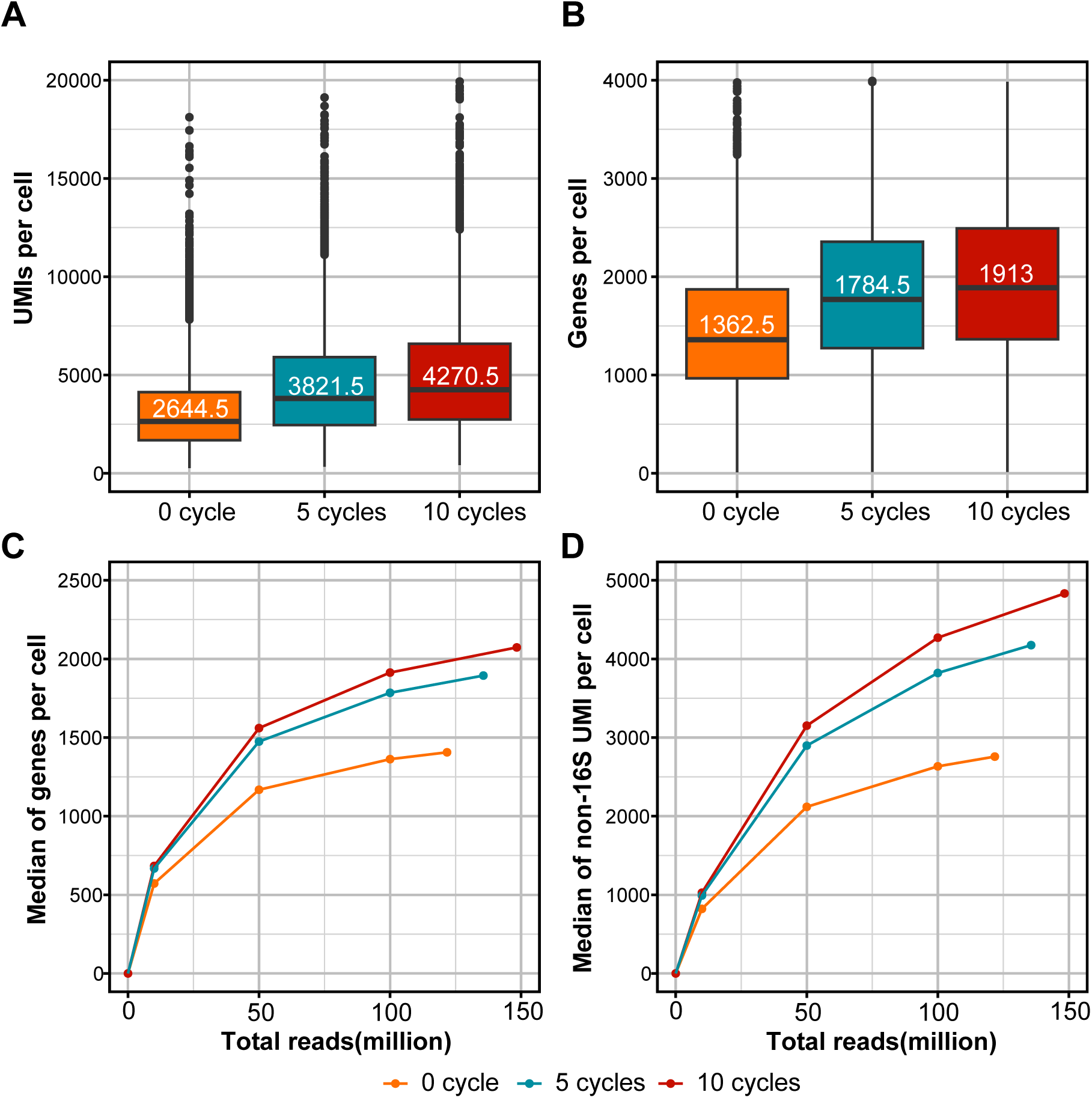
Post-CRISPR amplification increases library complexity. **A-B,** Boxplots showing the UMI (**A**) and genes per cell (**B**) among the libraries with different numbers of PCR cycles post-CRISPR. The medians are labeled in the 50th percentile in the box. **C-D,** Rarefaction analysis of library complexity comparing the libraries with different numbers of PCR cycles post-CRISPR, shown are the medians of genes (**C**) and non-16S UMIs per cell (**D**). Each dot represents a downsampled replicate with indicated total reads.

Since the extra PCR steps were necessary for improving library complexity, we asked whether the extra PCR was biased to enrich certain transcripts or cells that are abundant in the library. Therefore, we compared the numbers of genes and UMIs in shared cells in untreated and DASHed libraries. We found that the number of genes and non-16S UMIs in each cell with and without DASH treatment showed strong linear correlations, suggesting the relative UMI abundance across cells was maintained after DASH treatment (R^2^>0.97) (Figure 5A-B). Slopes greater than 1 in this analysis are indicative of a global increase in library complexity across cells. Moreover, we asked if these increases exist locally across different read depths. When we binned the cells into different groups based on the number of genes or UMIs, the levels of increase were consistent across different groups (Figure 5C-D). Together, these findings show that DASH increases cell complexity in an unbiased manner.

**Figure 5.**
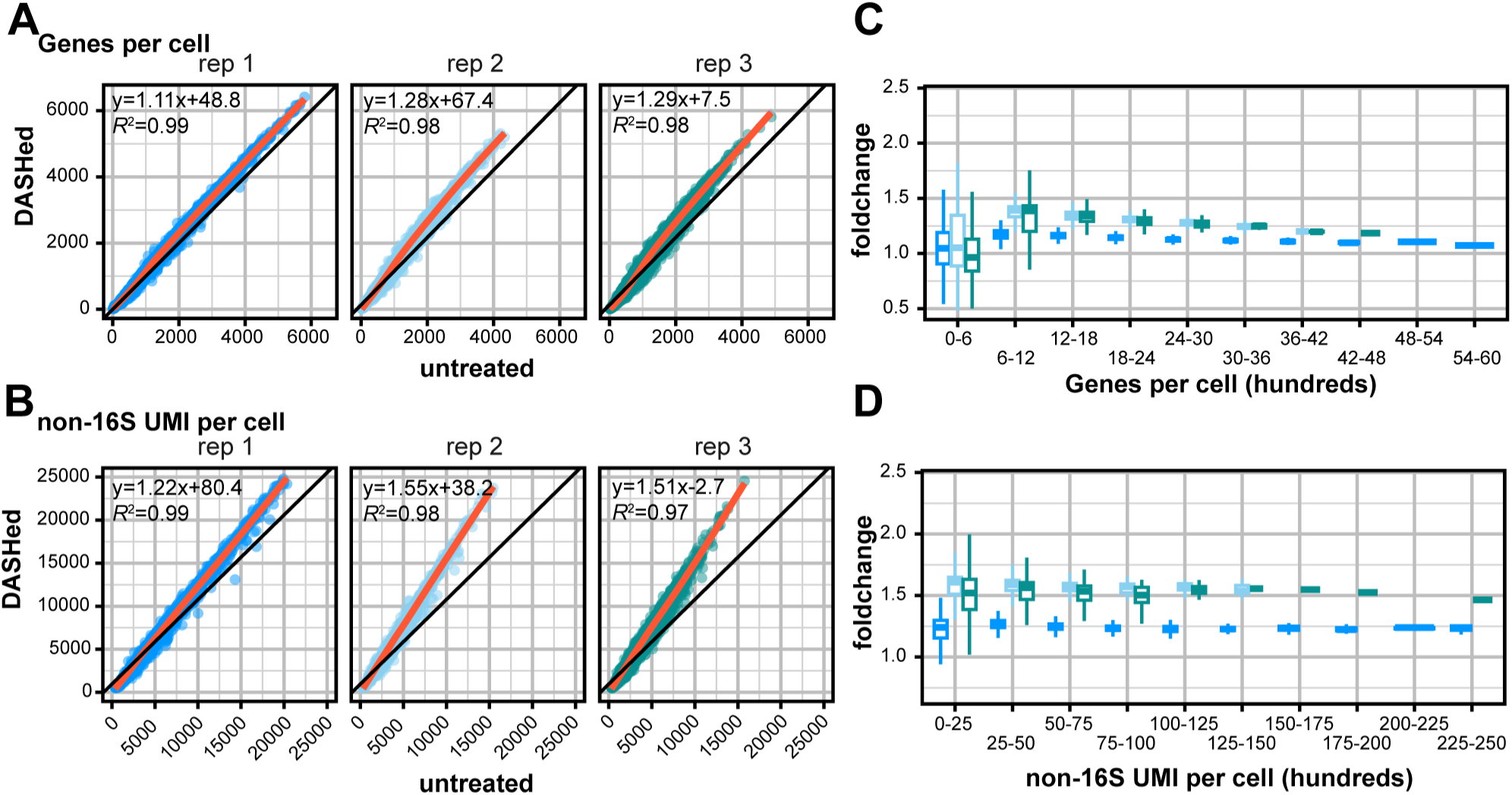
DASH treatment boosts library complexity independent of read depth. **A-B,** Comparison of genes per cell (**A**) and mRNA per cell (**B**) between untreated and DASHed libraries. Each dot represents a cell shared by untreated and DASHed. Orange lines are the linear regression models, and formula of linear regression and R-squared (R^2^) of the models are indicated in the top left. **C-D,** Boxplots show fold changes of DASHed vs. untreated in numbers of genes (**C**) and non-16S UMIs per cell (**D**) across ranges.

### Downstream single-cell analysis improves after DASH

By reducing contamination and ambient transcripts, the ultimate goal of DASH depletion is to reduce background and improve detection of rare cell types or more potential marker genes. To assess the benefit of DASH treatment as compared to the computational elimination of 16S, we removed 16S UMIs computationally from all libraries, and then pooled the replicates from either untreated or DASHed libraries. We used the shared nearest neighbor (SNN) method in Seurat to cluster the cells in both datasets separately (Supplemental Figure 4A-B). Clusters were annotated and aligned between two datasets based on previously described marker genes (19,20) (Supplemental Figure 4C), except that cluster 13 could not be annotated due to the high percentage of 16S sequence (Supplemental Figure 4D). Other clusters, with one exception, were present in both groups (Figure 6A-B). The unique cluster (C15) that appeared in the DASHed dataset represented *cathepsin*^+^ cells that were dispersed in the untreated dataset (Figure 6A), suggesting that the clustering is more sensitive due to the increased library complexity after DASH treatment.

**Figure 6.**
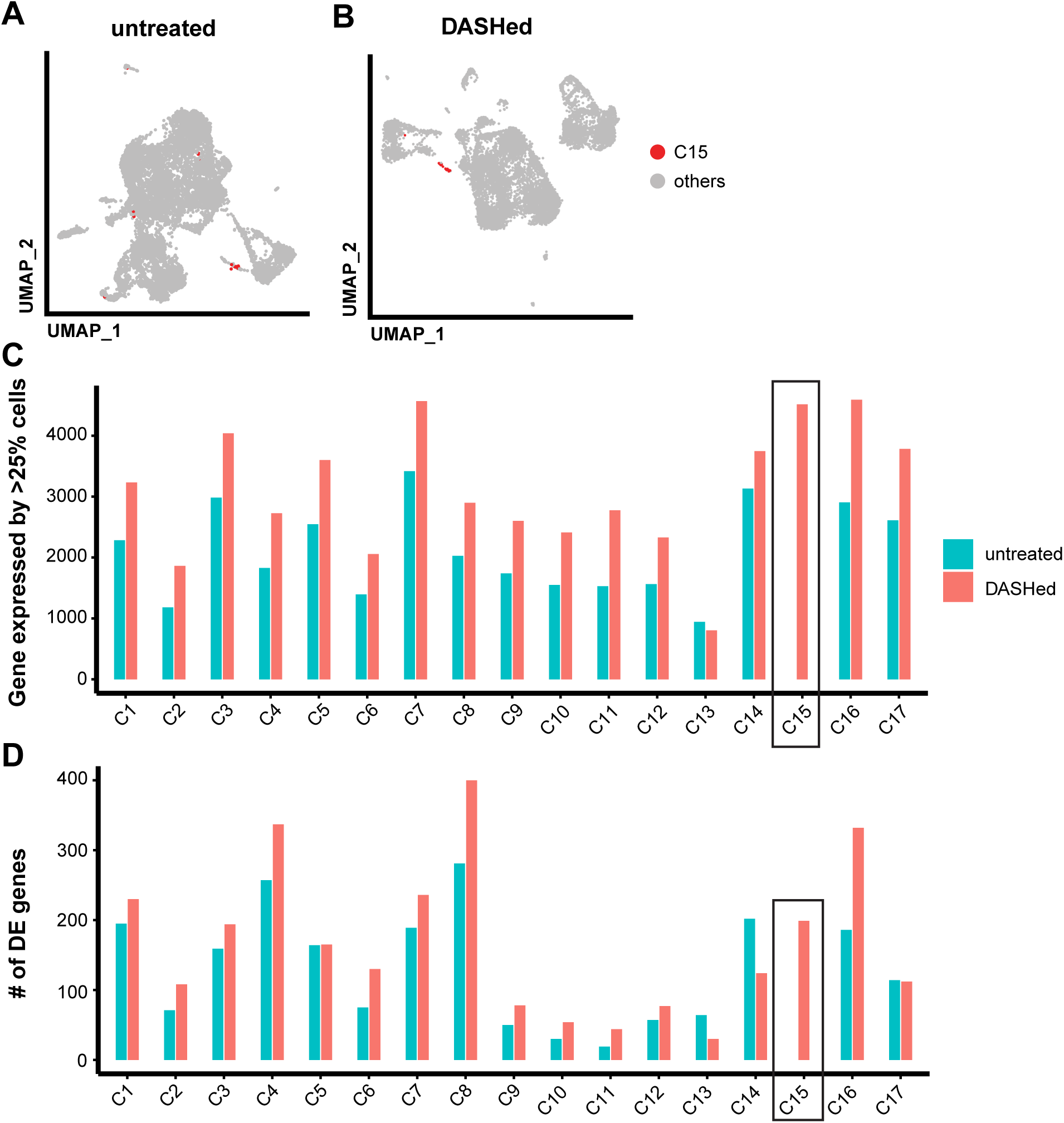
DASH treatment benefits clustering and differential expression analysis. **A-B,** UMAP plots of untreated (**A**) and DASHed (**B**) samples. Each dot represents a single cell. Cells belonging to cluster 15 (C15) in the DASHed sample are labeled in red. **C,** Bar plot shows numbers of genes that are expressed in at least 25% of cells in the same cluster. Black box indicates the cluster that is unique to the DASHed dataset. **D,** Bar plot shows numbers of differentially expressed (DE) genes across clusters, tested by Wilcoxon Rank Sum test and adjusted by Bonferroni correction. Black box indicates the cluster that is unique to the DASHed dataset.

Differential expression analysis in scRNA-seq often selects genes expressed in at least 10-25% of the cells within the cluster to obtain robust results (31). The number of genes expressed in at least 25% of cells of DASHed datasets (3090 genes on average) was higher than untreated datasets (2101 genes on average) across clusters, except for cluster 13, which has the highest enrichment for 16S (Figure 6C). To identify differentially expressed (DE) genes for each cluster, we used the Wilcoxon Rank Sum test. Of the 17 clusters, 14 had more DE genes in DASHed than untreated datasets (Figure 6D). We conclude that DASH treatment reduces the dropout rate of genes, enabling the clustering algorithm and differential expression analysis to perform better under the same parameters.

## Discussion

Sparsity of reads in single-cell sequencing is a technical challenge that arises from flooding of transcripts of housekeeping genes and ambient RNA that may swamp out detection of biologically informative genes (7,32). In this study, we analyzed several different scRNA-seq methodologies used in planarians and show that the single transcript for the planarian 16S rRNA accounts for 20-80% of all reads. To eliminate this contaminant, we integrated a DASH depletion step into the protocol for scRNA-seq library generation, and demonstrate that depletion of this sequence benefits overall library complexity. Removing the 16S transcript early during library preparation enhances the discovery of mRNAs that differentiate potential cell types, which we showed by performing a parallel analysis of untreated and depleted libraries. In conclusion, by eliminating unwanted reads, our approach improves detection of gene expression and economizes sequencing yield. Our approach is also customizable, and can be adapted to any system where similar contamination is evident.

### An unusually high fraction of 16S ribosomal RNA in planarian datasets

The prevalence of mitochondrial UMIs has been widely used as quality control for identifying healthy cells in human and mouse datasets, where acceptable maximum levels range from 5-10% (33,34). In the planarian *S. mediterranea*, mitochondrial 16S rRNA overflows RNA-seq experiments even after poly(A) enrichment (18). Following poly(A) enrichment, 16S still makes up 11-32% of total reads in bulk RNA-seq, and worsens in various single-cell methods, comprising 5-74% of total UMIs. While in most studies, the abundance of mitochondrial RNA is thought to arise from damaged cells, this is an unlikely source of 16S in planarians because most studies used either fresh or FACS-purified live cells (18,19,21). Ribosomal RNA transcribed in the nucleus is typically not polyadenylated, but reports have shown that both human and *Drosophila* mitochondrial 16S and 12S rRNA do get polyadenylated (29,35). Our coverage plots show that sequencing reads skew toward the 3’ end of 16S transcript, a trend resembling that of protein-coding genes. Alternatively, 3 poly(A) stretches in the middle of the 16S sequence might also contribute to its abundance, but this is unlikely because if reverse transcription initiated internally, reads corresponding to the 5’ end would be enriched over the 3’ end. Thus, we speculate that the planarian 16S might have an exceptionally long poly(A) tail, leading to significant capture in scRNA-seq, which remains to be tested experimentally. Overall, we capitalize on the overrepresentation of 16S to investigate the impact of ribodepletion on scRNA-seq.

### Alternative strategies for depleting abundant transcripts in single-cell RNA-seq

Recent advances in the ribodepletion of scRNA-seq have been shown to improve library complexity, but most of them require using new set-ups or novel reagents (8,10). Our DASH-mediated approach is straightforward to implement and highly customizable to any system (11). DASH has been shown to enhance library complexity in general but has been incorporated into single-cell methods in different ways (11,14,16,17). A common workflow of single-cell transcriptomes includes these steps in order: concomitant barcoding and reverse transcription, fragmentation of cDNA, and library indexing (3,4). Here, we digest the 16S rRNA before the fragmentation and indexing steps of the 10X Chromium protocol. To minimize PCR bias and loss of rare transcripts, we used 10 PCR cycles, 2 cycles fewer than recommended by 10X Genomics, to amplify cDNA before CRISPR/Cas9 digestion. After CRISPR digestion, we conduct a post-CRISPR amplification to enrich cDNA diversity, resulting in less than 0.5% of total UMIs belonging to 16S. Although other single-cell total transcriptome methods include CRISPR/Cas9-mediated depletion strategies to remove ribosomal sequences, the depletion is not as complete as what we observe here (10% of reads for Smart-seq-total (14) and 34% for scDASH (16) mapped to the target genes). More recently, a technique called scCLEAN targets 255 housekeeping and ribosomal genes for removal during library preparation. Even after depletion, reads mapping to these genes still constitute 8% of the total reads (17). The especially strong depletion that we observe here may result from several factors: (1) the contaminant in planarians is just a single transcript, and therefore easier to target and remove, (2) we used 30 sgRNAs to target one gene, compared to other approaches that use fewer sgRNAs per gene, and (3) eliminating target genes as early as possible in the library preparation may be beneficial.

### Impact of ribodepletion by DASH

While other studies have shown the overall benefit of DASH on library quality, we performed a parallel analysis of the same library before and after depletion to show the impact of DASH at single-cell resolution. This analysis reveals two key benefits of depletion. First, our data demonstrate that not depleting the 16S transcript leads to aberrant cell calling. The extra cells in untreated libraries are highly enriched in 16S UMIs, indicating that ambient RNA can falsify the cell calling process (36). Moreover, the distribution of transcripts during initial PCR steps required for library generation is distorted by their presence. Second, depletion of 16S does not appear to introduce bias in the analysis. We conclude this because the increase in library complexity and fold change of genes increased uniformly across cells with different UMIs. Furthermore, clustering analysis showed overall agreement between untreated and DASH-treated datasets and improvement in detecting differentially expressed genes by reducing the dropout rates. These findings are important because most scRNA-seq normalization uses a “size factor”, which equalizes the cell read depth (37). If the depletion happens disproportionately to the read depth across cells, normalization outcomes would be significantly altered by DASH. Future work using any depletion of a new panel of genes should look for any biased or off-target depletion.

In summary, we showcased the efficiency and robustness of DASH in scRNA-seq by demonstrating ribodepletion of 16S in planarians. The customizability of DASH will benefit any model organisms that may suffer from contamination of ambient RNA, or overabundance of irrelevant transcripts in important scRNA-seq experiments. The integration of DASH is a simple add-on to the current 10X protocol and therefore requires little expertise in developing new single-cell protocols. In addition, because the depletion can be done at any time, our protocol offers significant flexibility for sequencing after library generation.

## Conclusions

This study describes and benchmarks the ribodepletion of 16S rRNA by CRISPR-based treatment to improve scRNA-seq of planarians. Ribodepletion enhances the library complexity and performance of single-cell analysis. This demonstrates the benefit of ribodepletion in the cDNA library over *in silico* removal of ribosomal RNA.

## METHODS

### Worm care

*Schmidtea mediterranea* asexual clonal line CIW4 was raised in a recirculating water system supplemented with water containing Montjuïc salts (38,39). Animals were fed with beef liver and cleaned once a week. Animals were transferred to static culture containing 50 µg/mL gentamicin for at least a week prior to use.

### Cell sorting

For each biological replicate, 10 animals were dissociated into single-cell suspensions by dicing in CMFB buffer [calcium-magnesium-free solution with 1% BSA (400mg/L NaH_2_PO_4_, 800 mg/L NaCl, 1200 mg/L KCl, 800 mg/L NaHCO_3_, 240 mg/L glucose, 1% BSA, 15 mM HEPES, pH7.3)] and nutating for 2 hours at room temperature. Cells were centrifuged at 500g for 5 min, resuspended, and filtered through a 30µm cell strainer (BD Biosciences, 340628) to remove debris. The concentration of filtered cells was calculated using a TC20 automated cell counter (Bio-Rad). After centrifugation, cell concentration was adjusted to 100,000 cells/mL with staining buffer [CMFB containing DRAQ5 (5 µM) and Calcein-AM (0.4 µM)] and nutated at room temperature for 5 min. X1 cells were gated for vital 4N cells (DRAQ5^+^ Calcein-AM^+^) on a Sony MA900 Cell Sorter. 100,000 cells were sorted and diluted to a concentration of 1000 cells/µL for subsequent library preparation.

### sgRNA design and *in vitro* transcription

The detailed protocol for DASH is in Supplemental File 1. The 16S ribosomal RNA sequence was retrieved from the mitochondrial genome of *Schmidtea mediterranea* (NCBI accession number: NC_022448.1) and uploaded to IDT’s Alt-R Custom Cas9 crRNA Design Tool. 30 non-overlapping seed regions were selected from the output of the Design Tool to ensure complete digestion of the entire 905 bp transcript. The primer sequences are listed in Supplemental File 1. To synthesize T7-flanking templates for sgRNA, PCR reactions were assembled following the Phusion High-Fidelity (NEB) protocol with final concentrations of primers: one of the sgRNA primers (0.2µM), T7RevLong (0.2µM), T7FwdAmp (1µM), T7RevAmp (1µM). PCR reactions were carried out as follows: 98°C 30 sec, repeating the steps of 98°C for 10 sec, 51°C for 10 sec, 72°C for 10 sec 30 times, and then 72°C for 2min. PCR products were run on agarose gels to determine whether primers remained. If they did, the PCR products were gel purified. To synthesize sgRNA, the concentration of templates of each sgRNA was measured by nanodrop and pooled equivalently. In vitro transcription reactions were assembled as followed: sgRNA templates (4µg), 10X transcription buffer [0.1M MgCl_2_, 0.4 M Tris (pH 8.0), 0.1M DTT, 20 mM spermidine] (10µL), 25mM rNTPs (Promega) (8µL), T7 polymerase (generated in-house) (2µL), TIPP (NEB) (2µL), rRNAsin (Promega) (1µL) and nuclease-free water (adjusting the total volume to 100 µL). In vitro transcription reactions were incubated at 37°C overnight. The next day, 2 µL RQ1 DNase (Promega) was added to remove templates and incubated at 37°C for 20 min. To precipitate the sgRNAs, 250 µL ice-cold 100% ethanol was added to each reaction and incubated at −20°C for 1 hour. sgRNAs were pelleted by centrifugation at 4 for 2 minutes at 17,000g, and the supernatant was removed. To wash, 250 µL 70% ice-cold ethanol was added, followed by centrifugation for 2 minutes at 17,000g twice. The sgRNAs were resuspended in 10 µL nuclease-free water.

### 10X single-cell library preparation and DASH

Sorted cells were counted and checked viability on a Countess 3 Automated Cell Counter (ThermoFisher) with Trypan blue (0.4%) staining. Sorted cells showing >85% viability were used for 10X single-cell library preparation. To aim for recovery of 5000 cells after sequencing, 8250 cells were loaded onto the 10X Genomics Chromium Controller for subsequent library preparation using Chromium Next GEM Single Cell 3 Reagent Kit v3.1. Samples were amplified with 10 PCR cycles with cDNA primers (R1+TSO) after clean-up. For DASH, 29 µL CRISPR master mix [NEBuffer 3.1 (3µL), 20 μM sgRNAs (1µL, final 0.66μM), 1 µM Cas9 Nuclease (NEB, M0386S) (2µL, 66nM), nuclease-free water (23 µL)] was mixed and pre-incubated at 37 for 10 min. Then, 1µL (1-10 ng/µL) of cDNA was added to the master mix and incubated at 37 overnight. After CRISPR treatment, the cDNA was cleaned up and eluted into 15 µL using AMPure beads (Beckman) following the manufacturer’s protocol. Then, the cDNA was diluted to 30 µL and amplified with cDNA primers (R1+TSO) again with 10 cycles unless otherwise specified in this study. After PCR amplification, the cDNA was processed as specified in the 10X Genomics protocol for enzymatic fragmentation and indexing. Three biological replicates were used in this study. Libraries were pooled and sequenced using the NextSeq 2000 platform (Illumina). To assess whether 16S cDNA was removed, we ran individual samples on the Fragment Analyzer (Agilent).

### Library quality metrics for published datasets

Parallel-fastq-dump (0.6.7) was performed to retrieve fastq files from NCBI, and the SRA accession numbers were in Supplemental Table 1. All the preprocessing and alignment in Figure 1 use Drop-seq tools (2.3.0) except Smart-seq2 (3). Smart-seq2 method doesn’t have unique molecular modifiers (UMIs), so Smart-seq2 dataset was processed differently (40). Reads of Smart-seq2 were aligned to a customized genome file containing both the chromosomal-level (Smed_chr_ref_v1) (27) and mitochondrial genomes of *Schmidtea mediterranea* (NCBI accession number: NC_022448.1) using STAR (2.7.10) (41), and reads mapped to exon regions annotated by SMESG gene model were extracted and segregated into a gene expression matrix. In other single-cell RNA-seq libraries, reads were aligned to the same genome file described above using Drop-seq tools. Cell barcodes and unique molecular identifiers (UMIs) were extracted and tagged to the reads in BAM format and further segregated into a gene expression matrix. To calculate library quality metrics, ‘CreateSeuratObject’ function of Seurat (4.3.0) in R(4.2.0) was used to import the count matrix from Drop-seq tools or Smart-seq2 pipeline, which calculates total UMI counts and the number of genes expressed associated with each cell barcode (31). Further, custom R codes were used to extract 16S UMIs and non-16S UMIs (total UMIs - 16S UMIs). The workflow was adapted from TAR-scRNA-seq and compiled in snakemake (7.18.2) (42).

### Single-cell analysis for untreated and DASHed libraries

Figure 2B demonstrates the analysis workflow. The untreated and DASHed libraries from three biological replicates were processed by Cell Ranger (6.1.2) (4). The lists of cell barcodes were retrieved from both libraries and split into “untreated-specific”, “shared” and “DASHed-specific” categories (Supplemental Figure 3). The percentage of 16S UMIs was calculated for each category. For the rest of the analysis, only “shared” cells were used.

For rarefaction analysis, we downsampled the datasets into 10, 50, and 100 million reads for replicate pairs 2 and 3, and 150 and 200 million reads for replicate pair 1. Downsampled datasets were processed independently in CellRanger.

For clustering analysis, the shared cells were further selected if the cells had (1) more than 200 genes and (2) *piwi-1* expression ≥ 2.5 [ln(UMI-per-10,000 + 1)] to remove cells with low complexity and non-stem cells. If a cell didn’t meet the criteria in the untreated dataset, the same cells would be also eliminated in the DASHed dataset. Subsequently, all three replicates of either untreated or DASHed libraries were pooled separately. The libraries were normalized and scaled in Seurat and then integrated by Harmony (0.1.1) (43). The first fifty Harmony coordinates were used to calculate UMAP embedding. The clustering used the first twenty Harmony coordinates with a resolution of 0.5 for the FindClusters function, resulting in 16 clusters in untreated and 17 clusters in DASHed datasets (31). The list of markers from Zeng et al. (2018) was used to manually annotate and align the clusters (20). To detect differentially expressed genes, Wilcoxon Rank Sum adjusted by Bonferroni correction was used to compare untreated and DASHed samples (Figure 2E) or across clusters (Figure 6C-D). Differentially expressed genes (1) had an adjusted p-value <0.05, (2) were expressed by at least 25% of cells within the clusters and (3) log2 fold change >0.25 compared to other clusters.

## Declarations

### Ethics approval and consent to participate

Not applicable

## Consent for publication

Not applicable

## Availability of data and materials

All data is available upon request. scRNA-seq data have been deposited into NCBI with Accession number GSE231548. Code for generating the figures can be found on github https://github.com/kw572/DASH_figures.

## Competing interests

The authors declare that they have no competing interests.

## Funding

This work was funded by a National Institutes of Health grant R01GM139933 (to CEA), and Cornell University Mong Fellowship (to K-TW).

## Authors’ contributions

K-TW and CEA conceived and designed the study and drafted the manuscript; K-TW acquired data and performed computation analyses; CEA supervised the study. All authors read and approved the final manuscript.

## Supporting information

Supplemental Table and protocol

## Acknowledgments

We thank the Cornell University Biotechnology Resource Center’s Flow Cytometry (RRID:SCR_021740), Imaging (RRID:SCR_021741), and Genomics (RRID:SCR_021727) cores. We would like to thank Josien van Wolfswinkel for the suggestion to analyze dropout rates. We would also like to thank members of the Adler laboratory, David McKellar, Charles Danko, Leslie Babonis, and Bhargav Sanketi for comments on the manuscript.

---

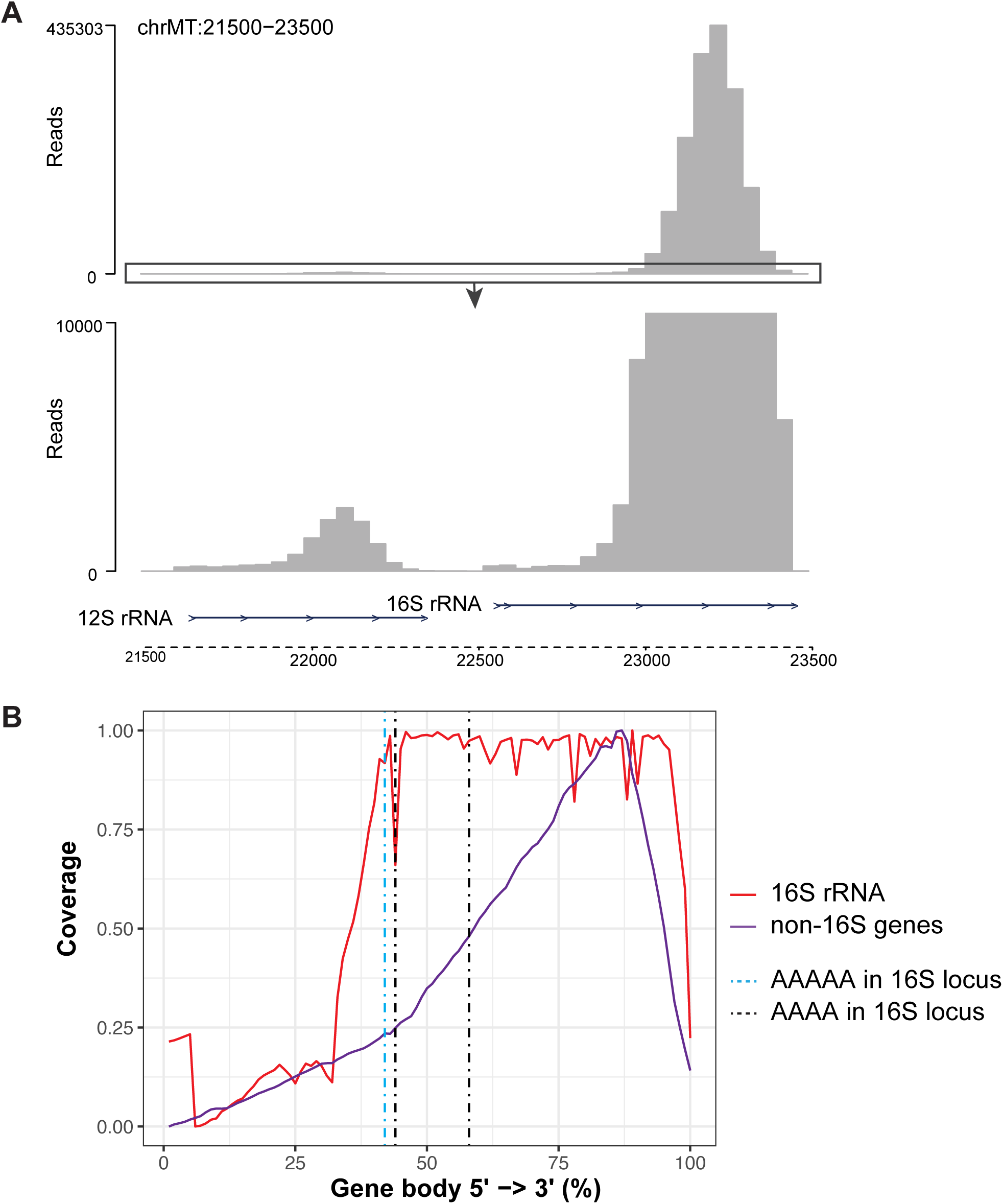

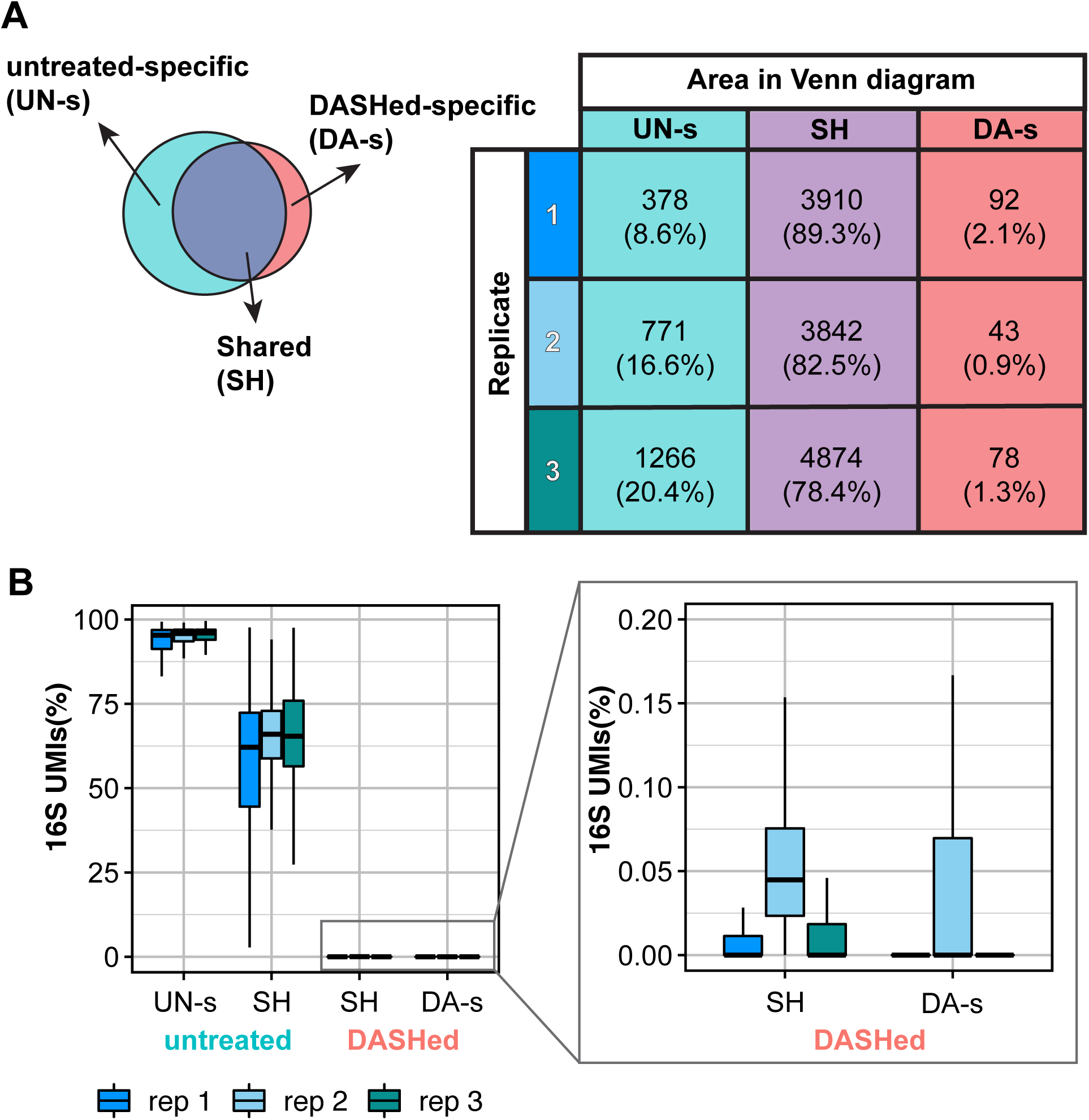

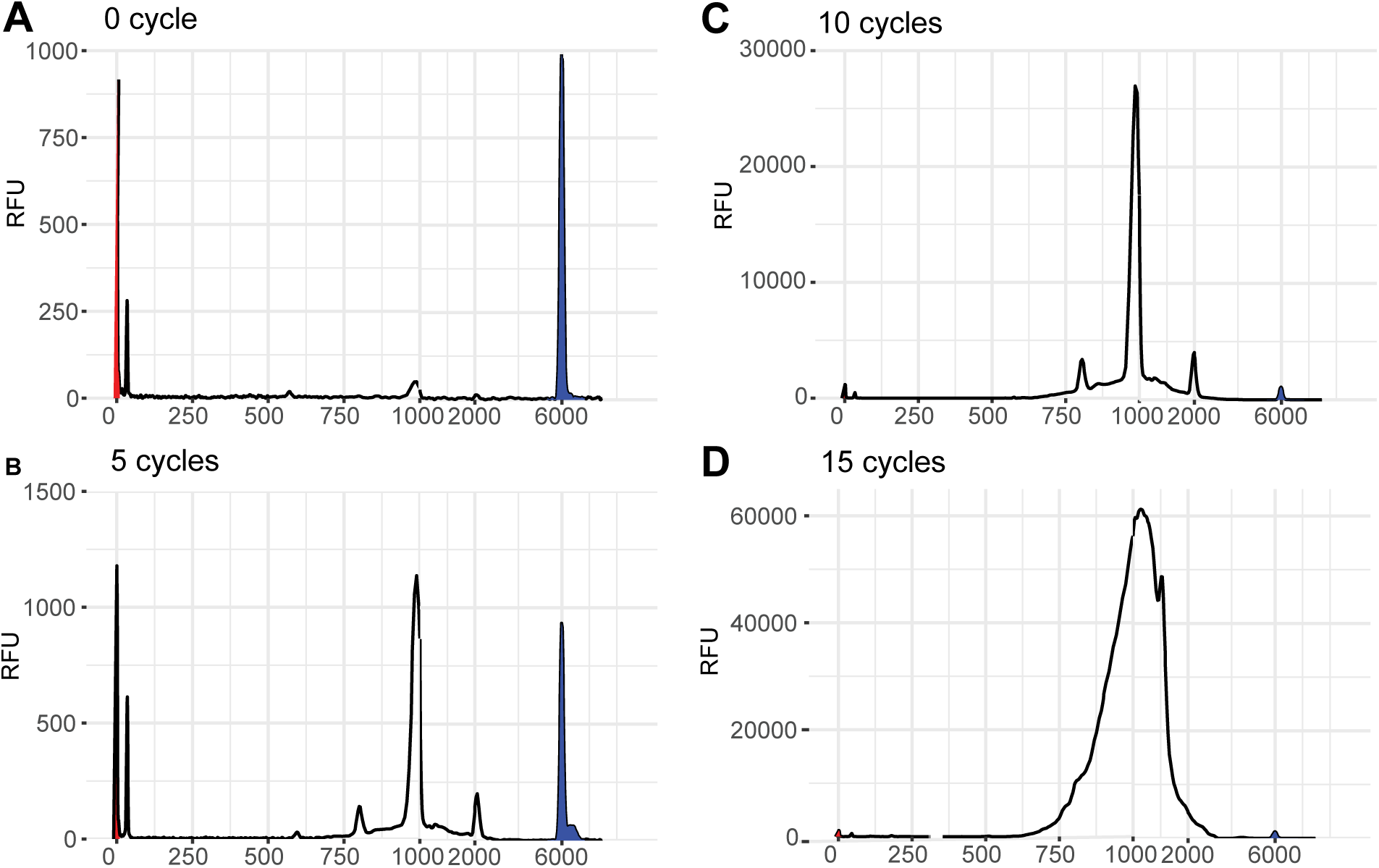

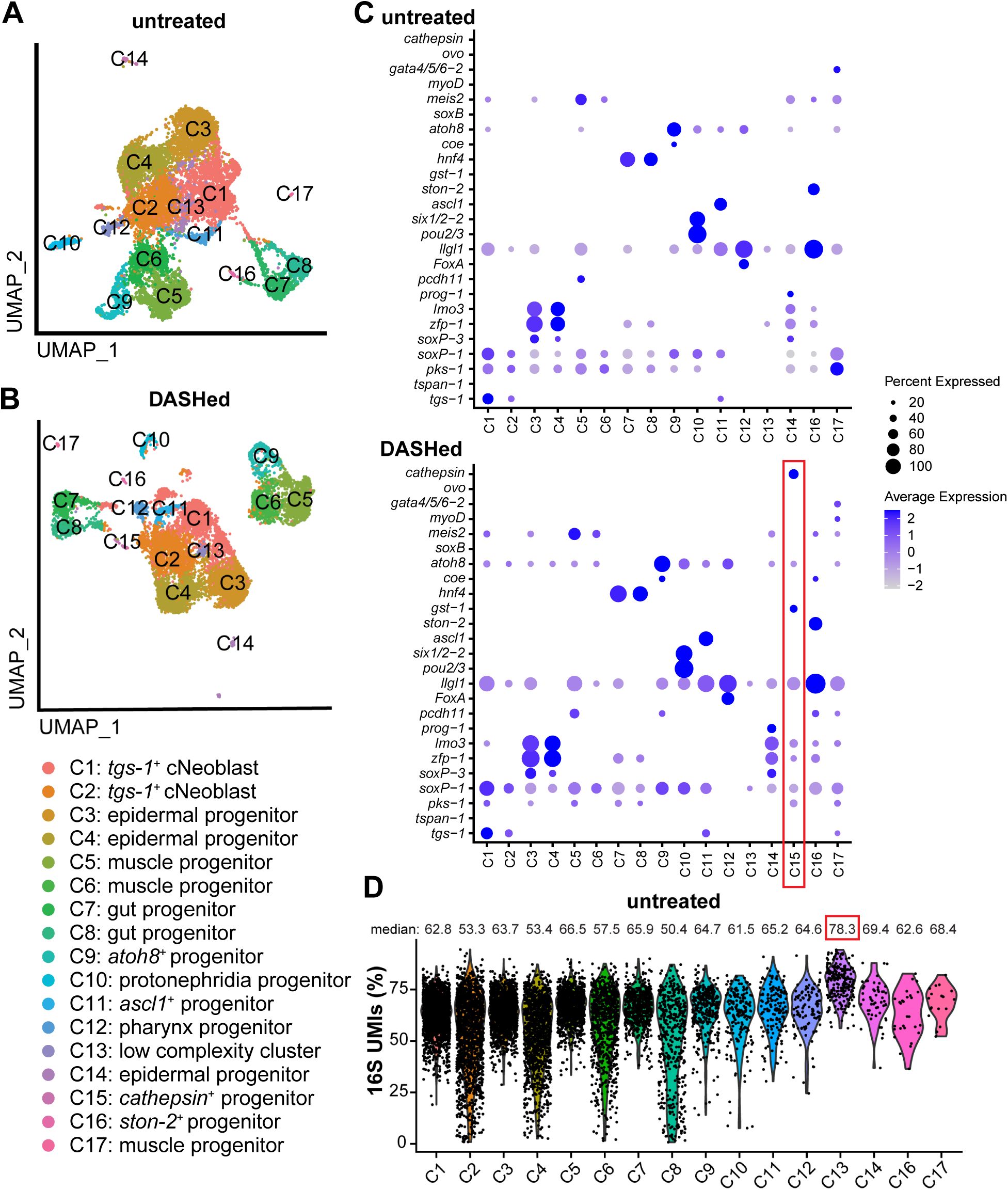

## Notes

### Competing Interest Statement

The authors have declared no competing interest.

### Summary of Updates

We corrected an error in the concentration of sgRNAs used for DASH treatment in the protocol and methods.

