## Supplemental Table and protocol for "CRISPR/Cas9-based depletion of 16S ribosomal RNA improves library complexity of single-cell RNA-sequencing"

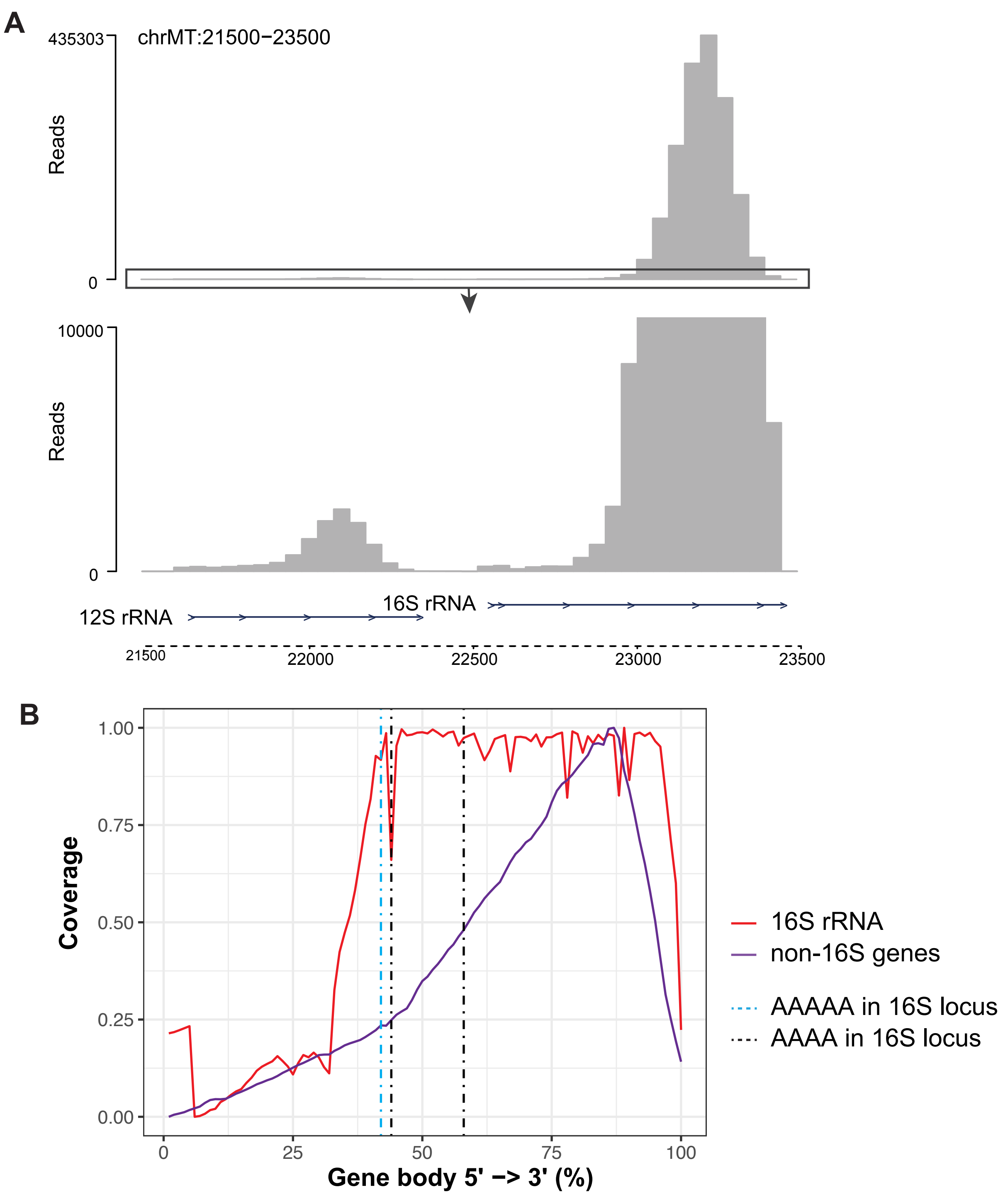


**Supplementary figure 1. 16S reads are enriched at the 3' end of the locus.**
**A,** Genomic tracks showing the raw read depth surrounding 12S and 16S rRNA loci, The bottom track shows 0-10000 to highlight the 12S locus. Arrows show the orientations of gene loci.
**B,** Gene body coverage plot showing the coverage of 16S rRNA locus (red) and average coverage of all non-16S gene loci (purple) in planarians. The coverage is retrieved from the untreated library, replicate 2. x-axis represents the percentage of the gene body, from 0% (5' end) to 100% (3' end). y-axis represents the relative coverage of reads that map to each part of the gene body. Dashed lines indicate the positions of A stretches in 16S locus.

A

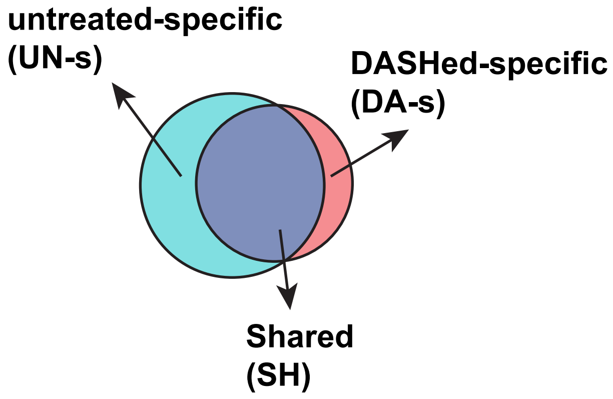


| | | Area in Venn diagram | | |
|---|---|---|---|---|
| | | UN-s | SH | DA-s |
| Replicate | 1 | 378 (8.6%) | 3910 (89.3%) | 92 (2.1%) |
| | 2 | 771 (16.6%) | 3842 (82.5%) | 43 (0.9%) |
| | 3 | 1266 (20.4%) | 4874 (78.4%) | 78 (1.3%) |

B

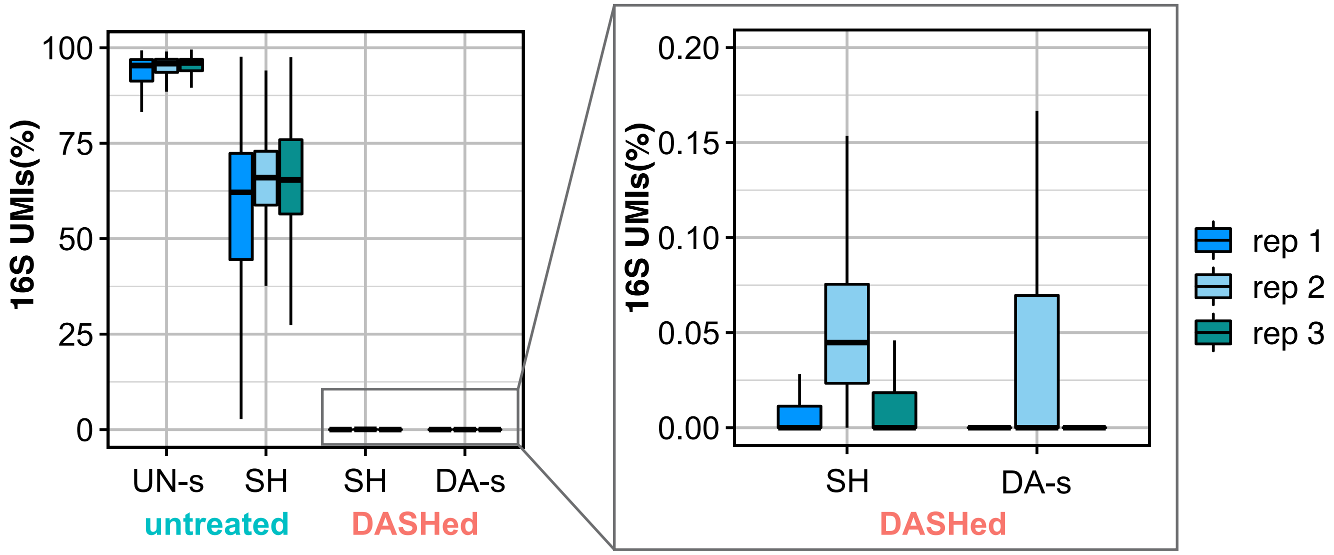


**Supplementary figure 2. Overrepresentation of 16S UMI causes aberrant cell calling.**
**A,** Venn diagrams show the cell barcodes shared or specific to “untreated” or “DASHed” across biological replicates. Table shows the number of cell barcodes in each area and replicate. The percentages show the proportion of the numbers over the total cell numbers within the replicate.
**B,** Box plots depict the percentage of 16S UMI from the cells of indicated area. Enlarged box plots from B show cells from “DASHed” libraries.

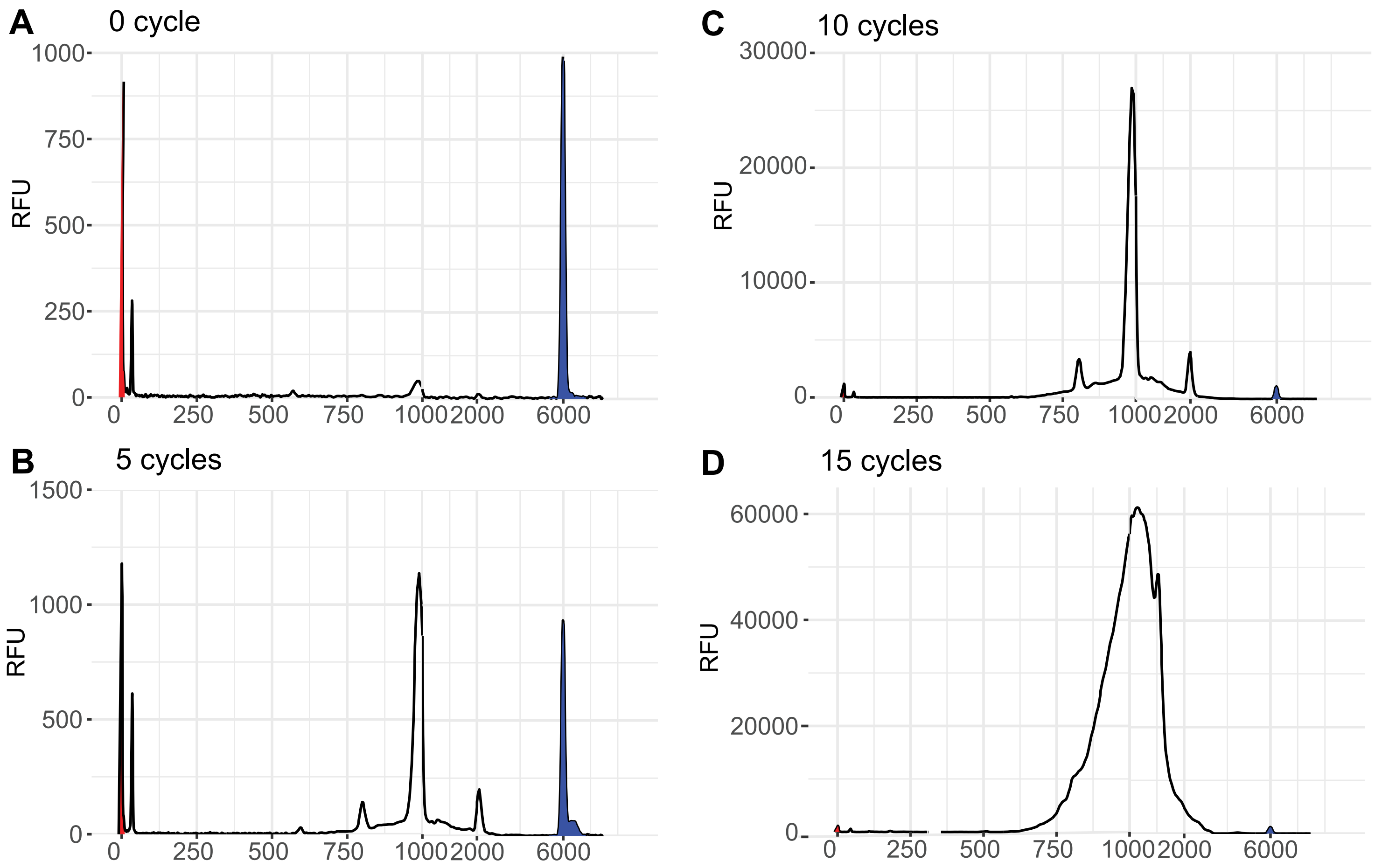


**Supplementary figure 3. 10 PCR cycles post-CRISPR is optimal.**
Fragment analysis of cDNA with 0 (A), 5 (B), 10 (C) and 15 (D) cycles post-CRISPR.
x-axis shows fragment size in base pairs (bp), and y-axis is relative fluorescence units (RFU). Red peak marks the lower marker (1bp) and blue peak marks the upper marker (6000bp).

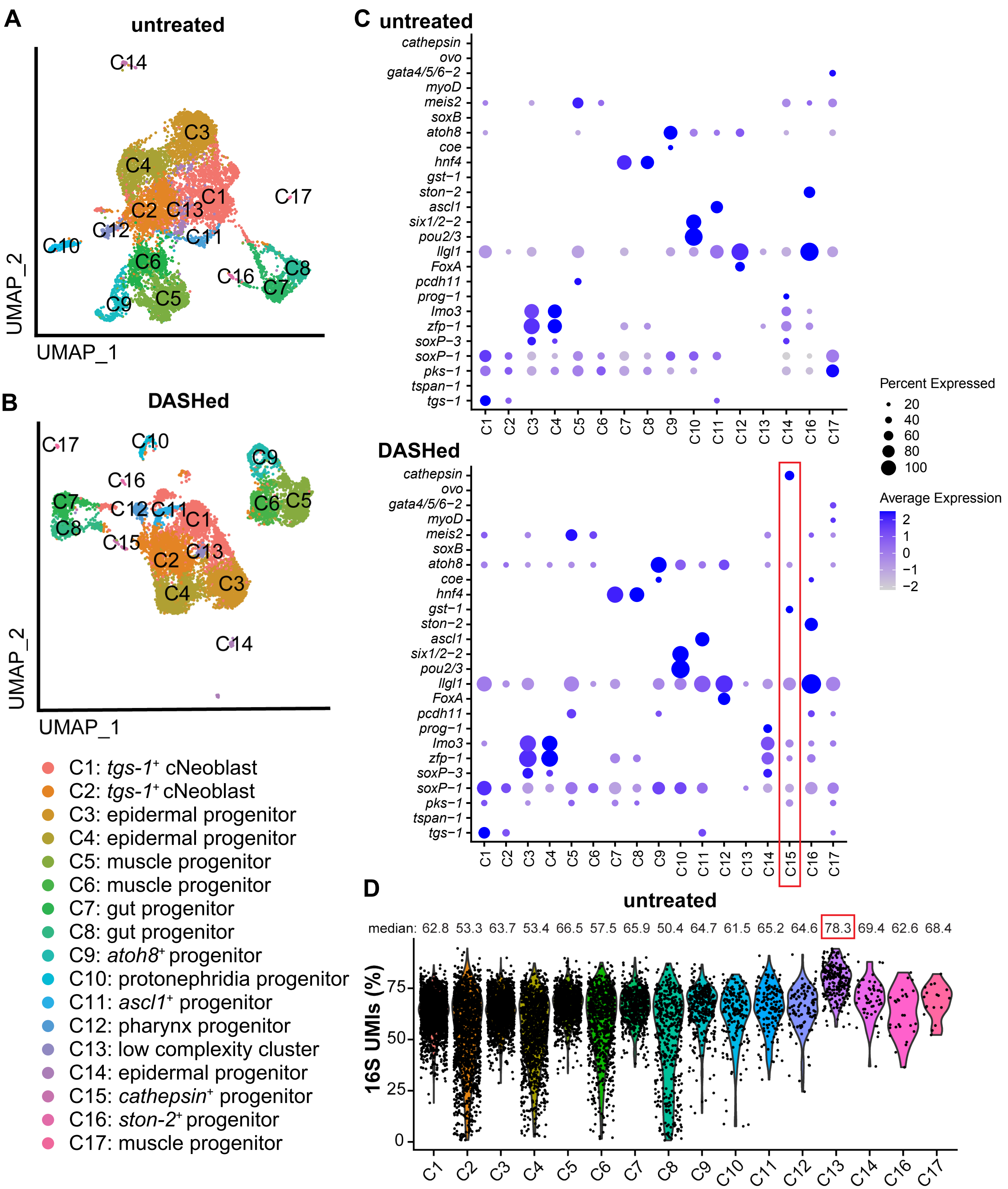


**Supplementary figure 4. DASH treatment benefits clustering and differential expression analysis.**
**A-B,** UMAP plots of untreated (A) and DASHed (B) samples. Each dots represent a single cell. Dots are color-coded by clusters.
**C,** Dot plot of the marker gene expression across annotated clusters. The size of the dots represents the percentages of cells within each cluster that express the marker gene. The color gradient represents the expression level. Red box shows the cluster unique to DASHed dataset.
**D,** Violin plot shows the percentage of 16S UMIs in each cluster, where C13 has the highest (red box). Each dot represents a single cell.

**Table 1: Quality metrics of planarian single cell datasets**

| dataset | Median of genes | Median of UMIs | Mean of genes | Mean of UMIs | 16S% | 12S% | mito% | Total Reads | Uniquely mapped reads % | % of reads mapped to multiple loci | % of reads unmapped: too short | SRA files |
|---|---|---|---|---|---|---|---|---|---|---|---|---|
| ACME | 313.5 | 654.5 | 421.75 | 898.88 | 25.24% | 0.50% | 1.32% | 99,760,985 | 21.90% | 21.10% | 53.94% | SRR11768230 |
| Fincher.1 | 627.5 | 1447 | 870.32 | 2418.39 | 21.45% | 0% | 0.67% | 21,927,812 | 65.08% | 10.24% | 21.13% | SRR6829380 |
| Fincher.2 | 645.5 | 1588 | 868.04 | 2473.85 | 25.36% | 0% | 0.78% | 50,022,789 | 60.08% | 9.50% | 25.55% | SRR6829381 |
| Fincher.3 | 657.5 | 1486 | 872.37 | 2433.80 | 25.56% | 0% | 0.89% | 99,120,749 | 63.57% | 9.85% | 21.94% | SRR6829382 |
| Fincher.4 | 936 | 2381 | 1031.81 | 3007.32 | 26.64% | 0.03% | 1.14% | 487,831,951 | 69.84% | 10.95% | 14.87% | SRR6829389;<br>SRR6829390;<br>SRR6829391;<br>SRR6829392 |
| Fincher.5 | 886 | 2373 | 972.57 | 2866.42 | 32.87% | 0.04% | 1.61% | 332,546,131 | 68.65% | 9.71% | 18.33% | SRR6829395;<br>SRR6829396;<br>SRR6829397 |
| Plass | 349 | 1365 | 456.76 | 1801.93 | 52.05% | 0.38% | 4.44% | 152,686,345 | 75.57% | 7.85% | 14.06% | SRR6020457 |
| Scimone | 1648 | 14338 | 1658.40 | 14723.31 | 74.37% | 0.74% | 2.24% | 246,559,964 | 74.28% | 7.07% | 16.63% | SRR15016138 |
| SmartSeq2 | 1623 | 706915 | 1821.77 | 811164.43 | 39.18% | 0.12% | 5% | 1,473,623 | 32.64% | 35.20% | 27.18% | GEO:GSE79866 |
| SplitSeq.1 | 406 | 647 | 475.79 | 807.41 | 5.15% | 1.26% | 0.54% | 52,438,770 | 47.45% | 39.08% | 4.86% | SRR11276863 |
| SplitSeq.2 | 891.5 | 1524 | 1011.77 | 1821.92 | 4.54% | 1.30% | 0.87% | 96,156,650 | 48.35% | 39.11% | 5.29% | SRR11276865 |
| SplitSeq.3 | 348 | 613.5 | 433.34 | 811.40 | 8.36% | 3.58% | 1.41% | 76,997,859 | 44.66% | 43.20% | 4.96% | SRR11276867 |
| Zeng | 2481 | 14358 | 2463.81 | 15166.24 | 60.58% | 0.26% | 3.25% | 562,100,456 | 81.59% | 4.62% | 13.69% | SRR6363908 |

### Single-cell DASH with 10X single-cell sequencing

#### Introduction

DASH effectively depletes unwanted cDNA sequences in bulk RNA-seq libraries with minimal input (1ng) by CRISPR-Cas9 cleavage. DASH has also been applied to several single-cell RNA-seq protocols[1–3]. Here, we demonstrate DASH is also adaptable to 10X single-cell sequencing pipeline and effectively depletes planarian 16S ribosomal RNA that dominates planarian single-cell transcriptomes. This depletion of 16S rRNA saves cost, enhances library complexity, and does not compromise downstream analysis.

#### Prepare sgRNAs[4]

1. The gRNAs are designed to target 16S in a non-overlapping manner by IDT guide RNA design tool.

| | |
|---|---|
| T7RevLong | AAAAAAAGCACCGACTCGGTGCCACTTTTTCAAGTTGATAACGGACTAGCCTTATTTAAACTTGCTATGCTGTTTCCAGCATAGCTCTTA |
| T7FwdAmp | TAATACGACTCACTATAG |
| T7RevAmp | AAAAAAAGCACCGACTCGGTGC |
| 16S_1 | TAATACGACTCACTATAGATATTTTCCTTTTTGTATTAGTTTAAGAGCTATGCTGGAAAC |
| 16S_2 | TAATACGACTCACTATAGCTTTTTGTATTACGGTTTGTGTTTAAGAGCTATGCTGGAAAC |
| 16S_3 | TAATACGACTCACTATAGTATGTTTTTATTCTCTCGAAGTTTAAGAGCTATGCTGGAAAC |
| 16S_4 | TAATACGACTCACTATAGGTGTTTTGTTTTATTTTCTTGTTTAAGAGCTATGCTGGAAAC |
| 16S_5 | TAATACGACTCACTATAGTGATAGTAGTGATATGTTTTGTTTAAGAGCTATGCTGGAAAC |
| 16S_6 | TAATACGACTCACTATAGTCTAGTTTTTAATTGTCTTTGTTTAAGAGCTATGCTGGAAAC |
| 16S_7 | TAATACGACTCACTATAGTAGTCGTTTGTTTGCTTATAGTTTAAGAGCTATGCTGGAAAC |
| 16S_8 | TAATACGACTCACTATAGATTTAACTTATCAGTGAGCAGTTTAAGAGCTATGCTGGAAAC |
| 16S_9 | TAATACGACTCACTATAGGCAGTACTTTGACTGTACGAGTTTAAGAGCTATGCTGGAAAC |
| 16S_10 | TAATACGACTCACTATAGTAATTCTAGAATTGTTTGAAGTTTAAGAGCTATGCTGGAAAC |
| 16S_11 | TAATACGACTCACTATAGAAAGTATAAATTAAAAAACAGTTTAAGAGCTATGCTGGAAAC |
| 16S_12 | TAATACGACTCACTATAGAGATACAGTTTGTTATTTCAGTTTAAGAGCTATGCTGGAAA |
| 16S_13 | TAATACGACTCACTATAGCTAGAGAGTTTTTAACTTAGGTTTAAGAGCTATGCTGGAAAC |
| 16S_14 | TAATACGACTCACTATAGTTCTTACTTTAGTTATTTGTGTTTAAGAGCTATGCTGGAAAC |
| 16S_15 | TAATACGACTCACTATAGTTAGTTATTTGTTGGGGTAAGTTTAAGAGCTATGCTGGAAAC |
| 16S_16 | TAATACGACTCACTATAGATTACATTATGAACTTCCTTGTTTAAGAGCTATGCTGGAAAC |
| 16S_17 | TAATACGACTCACTATAGTGAACTTCCTTAGGGATAACGTTTAAGAGCTATGCTGGAAAC |

| | |
|---|---|
| 16S_18 | TAATACGACTCACTATAGCTATATCCCTGTTATCCCTAGTTTAAGAGCTATGCTGGAAAC |
| 16S_19 | TAATACGACTCACTATAGATAACAGGGATATAGAATCTGTTTAAGAGCTATGCTGGAAAC |
| 16S_20 | TAATACGACTCACTATAGACAGGGATATAGAATCTTGGGTTTAAGAGCTATGCTGGAAAC |
| 16S_21 | TAATACGACTCACTATAGTTAACTACAATTCAACATCGGTTTAAGAGCTATGCTGGAAAC |
| 16S_22 | TAATACGACTCACTATAGGTTGAATTGTAGTTAAAGATGTTTAAGAGCTATGCTGGAAAC |
| 16S_23 | TAATACGACTCACTATAGGTAGTTAAAGATAGGTGTAGGTTTAAGAGCTATGCTGGAAAC |
| 16S_24 | TAATACGACTCACTATAGGTCGAACAGACTAAAAGTAAGTTTAAGAGCTATGCTGGAAAC |
| 16S_25 | TAATACGACTCACTATAGAGTTTAGACCGATGTGAATCGTTTAAGAGCTATGCTGGAAAC |
| 16S_26 | TAATACGACTCACTATAGTAGACCGATGTGAATCAGGTGTTTAAGAGCTATGCTGGAAAC |
| 16S_27 | TAATACGACTCACTATAGAAAACCAACCTGATTCACATGTTTAAGAGCTATGCTGGAAAC |
| 16S_28 | TAATACGACTCACTATAGCGTACAATGGACAGAAATTCGTTTAAGAGCTATGCTGGAAAC |
| 16S_29 | TAATACGACTCACTATAGGTCCATTGTACGAAAGGAATGTTTAAGAGCTATGCTGGAAAC |
| 16S_30 | TAATACGACTCACTATAGATCCAATTCCTTTCGTACAAGTTTAAGAGCTATGCTGGAAAC |

2. To synthesize gRNA templates, make PCR reactions as below:

| Component | Volume |
|---|---|
| Nuclease-free Water | 10.6 µl |
| 5x Phusion HF buffer | 4 µl |
| 10mM dNTPs | 0.4 µl |
| *16S_number | 0.4 µl |
| T7RevLong | 0.4 µl |
| T7FwdAmp | 2 µl |
| T7RevAmp | 2 µl |
| Phusion HF DNA polymerase | 0.2 µl |

| | |
|---|---|
| Total | 20 µl |

* 16S_number refers to each oligo name starting with 16S

3. Run PCR as below:

| | | |
|---|---|---|
| Step1 | 98℃ | 30 sec |
| Step2 | 98℃ | 10 sec |
| Step3 | 51℃ | 10 sec |
| Step4 | 72℃ | 10 sec<br>*30 times step 2-4 |
| Step5 | 72℃ | 2 min |
| Step6 | 12℃ | hold |

4. Run electrophoresis to make sure the PCR product is a single band without excessive primers
5. No PCR cleanup necessary. Otherwise, do gel purification.
6. Measure the concentration by nanodrop
7. Pool all the DNA templates equivalently
8. In vitro transcription
   Example:

| | |
|---|---|
| sgRNA templates | 4 µg |
| 10X buffer | 10 µl |
| rNTP 25mM | 8 µl |
| T7 polymerase | 2 µl |
| TIPP | 2 µl |
| rRNAsin | 1 µl |
| Nuclease-free water | Bring to 100 µl |

9. Add 1uL RQ1 RNAse-free DNAse, incubate RT for 20 minutes
10. Add 250uL 100% Ethanol
11. Incubate at -20℃ for 1 hour (can go overnight here)
12. Spin at max speed for 20 minutes at 4℃
13. Remove solvent from the pellet, and add 1mL ice-cold 70% ethanol
14. Vortex briefly and spin at max speed for 2 minutes at 4℃

15. Remove solvent from the pellet, and add 1mL ice-cold 70% ethanol
16. Vortex briefly and spin at max speed for 2 minutes at 4℃
17. Remove the solvent, and place the sample at 37℃ with the lid open until dry
18. Dissolve in the desired volume of the appropriate buffer. Suspending in 1/10th of the initial transcription reaction volume should yield around 2mg/mL sgRNAs

**Prepare 10X single cell libraries**[5]

1. Dissociate desired tissues into cell suspension
2. Stain and filter for FACS sorting
3. Collect targeted numbers of cells according to Chromium Single Cell 3' Reagent Kits User Guide (v3.1 Chemistry)
4. Resuspend into 1000 cells/μl in buffer recommended by Chromium Single Cell 3' Reagent Kits User Guide (v3.1 Chemistry). In this study, we use CMFB buffer [calcium-magnesium-free solution with 1% BSA (400mg/L NaH2PO4, 800 mg/L NaCl, 1200 mg/L KCl, 800 mg/L NaHCO3, 240 mg/L glucose, 1% BSA, 15 mM HEPES, pH7.3)]
5. Follow 10X chromium protocol and target 5000 cells/library until finishing Step 2 in Chromium Single Cell 3' Reagent Kits User Guide (v3.1 Chemistry)

**DASH treatment**[6]

1. After Step 2 in Chromium Single Cell 3' Reagent Kits User Guide (v3.1 Chemistry), the yield of cDNA will be 1-10ng/μl (from 3000-5000 X1 cells)
2. Assemble CRISPR master mix as below:

| Component | Volume | Final |
|---|---|---|
| Nuclease-free Water | 23 μl | |
| NEBuffer 3.1 | 3 μl | 1X |
| 20 μM sgRNA | 1 μl | 0.66 μM |
| 1 μM Cas9 Nuclease (M0386S) | 2 μl | 66 nM |
| Total | 29 μl | |

3. Pre-incubate for 10 min at 37℃
4. Add 1 μl of cDNA into CRISPR master mix
5. Incubate at 37℃ overnight

**cDNA purification**

1. Mix CRISPR reaction with 1.8X Ampure Beads (Beckman) with 10 times pipetting
2. Let stand for 5 min at room temperature
3. Place the tube on a magnetic stand for 3 min to separate the beads from the solution
4. Wash the beads with 80% freshly prepared EtOH 30 sec twice on the magnetic stand
5. Air dry for 3 min on the magnetic stand
6. Remove tubes from the magnetic stand and resuspend the beads with 15 μl Nuclease-free Water
7. Stand for 5 min at room temperature
8. Place the tube on a magnetic stand for 3 min to separate the beads from the solution
9. Transfer the solution into a new tube

**Library preparation**

1. Repeat Step 2.2 in Chromium Single Cell 3' Reagent Kits User Guide (v3.1 Chemistry) with 10 PCR cycles.
2. QC check on fragment analyzer
3. Proceed library preparation in Chromium Single Cell 3' Reagent Kits User Guide (v3.1 Chemistry)

**Expected results from fragment analyzer:**
**Before DASH:**

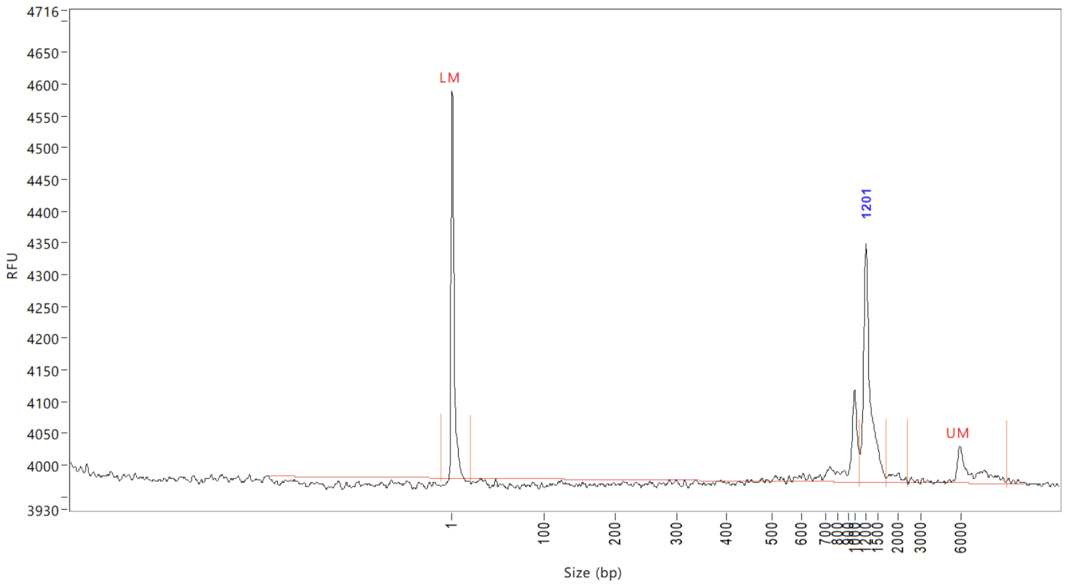


*Peak around 1200-1300 comes from 16S rRNA

**After DASH:**

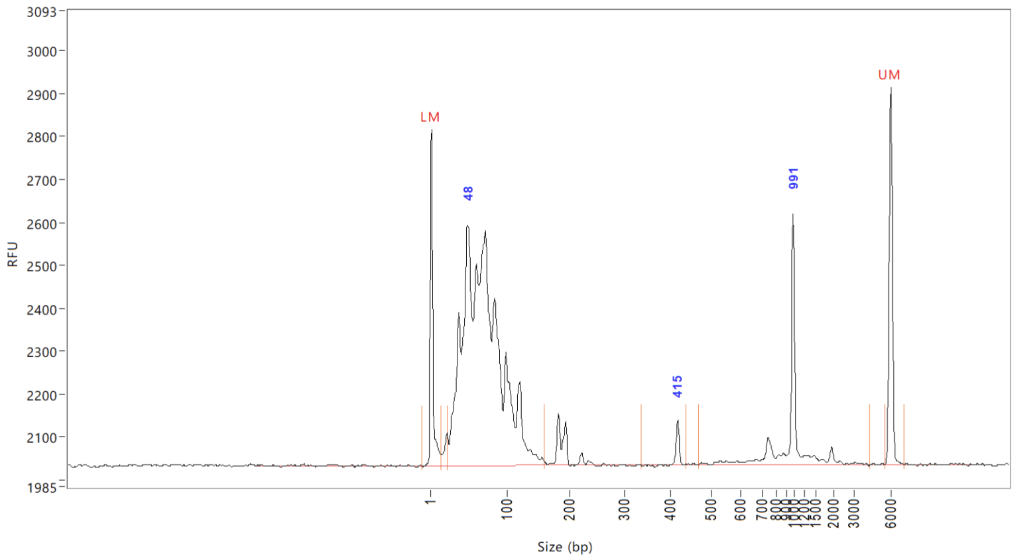


*The 16S rRNA peak should be gone, and the breakdown of 16S rRNA will be below 200 bp

**After DASH+10 PCRcycle:**

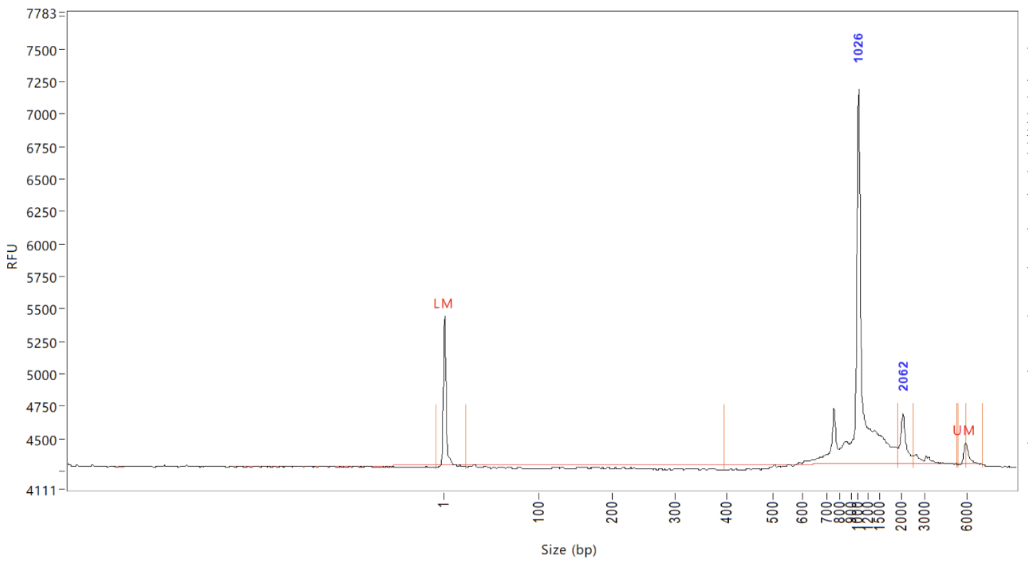


*Additional round of Step2.2 can get rid of 16S rRNA breakdown

**After fragmentation and indexing:**

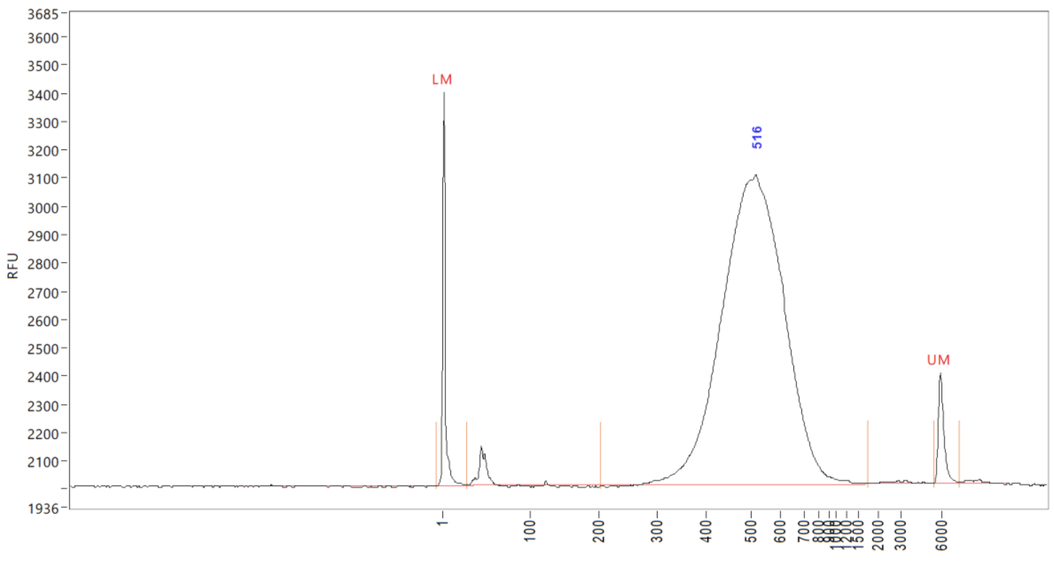

LM
516
UM
RFU
3685
3600
3500
3400
3300
3200
3100
3000
2900
2800
2700
2600
2500
2400
2300
2200
2100
1936
1
100
200
300
400
500
600
700
2000
3000
6000
